# Clonal differences underlie variable responses to sequential and prolonged treatment

**DOI:** 10.1101/2023.03.24.534152

**Authors:** Dylan L. Schaff, Aria J. Fasse, Phoebe E. White, Robert J. Vander Velde, Sydney M. Shaffer

**Affiliations:** Department of Bioengineering, School of Engineering and Applied Sciences, University of Pennsylvania, Philadelphia, PA, USA; Department of Chemistry, School of Arts and Sciences, University of Pennsylvania, Philadelphia, PA, USA; Department of Pathology, Perelman School of Medicine, University of Pennsylvania, Philadelphia, PA, USA

## Abstract

Cancer cells exhibit dramatic differences in gene expression at the single-cell level which can predict whether they become resistant to treatment. Treatment perpetuates this heterogeneity, resulting in a diversity of cell states among resistant clones. However, it remains unclear whether these differences lead to distinct responses when another treatment is applied or the same treatment is continued. In this study, we combined single-cell RNA-sequencing with barcoding to track resistant clones through prolonged and sequential treatments. We found that cells within the same clone have similar gene expression states after multiple rounds of treatment. Moreover, we demonstrated that individual clones have distinct and differing fates, including growth, survival, or death, when subjected to a second treatment or when the first treatment is continued. By identifying gene expression states that predict clone survival, this work provides a foundation for selecting optimal therapies that target the most aggressive resistant clones within a tumor.

## Introduction

Individual cancer cells frequently exhibit diverse responses to their environment and to treatments, resulting in some cells surviving while others die. These surviving cells can contribute to the development of resistant tumors, limiting the effectiveness of treatment. Although genetic differences often account for these varied responses^1–3^, non-genetic factors^4^, such as epigenetic state^5,6^, protein levels^7^, gene expression^8–11^, and cell cycle stage^12^, can also contribute to therapy resistance and survival in a hypoxic microenvironment^13,14^. Upon treatment, a subset of cancer cells survive and generate clones that are resistant to therapy. These resistant clones can exhibit significant interclonal heterogeneity, characterized by divergent gene expression states and invasive properties^9^. However, it remains uncertain how these heterogeneous resistant clones respond when subjected to a second treatment.

Second-line treatments are typically less effective compared to first-line treatments^15–18^. However, it is unclear how this occurs at the level of resistant clones or individual cells. Specifically, it is unknown whether all resistant clones respond uniformly to a second treatment or whether they exhibit different responses (Fig. 1A). Furthermore, when resistant clones undergo prolonged treatment with the same agent, it is unknown whether some clones grow more than others. While it is evident treatment can drive clones into diverse resistant states^9^, their phenotypic consequences have yet to be revealed.

**Figure 1:**
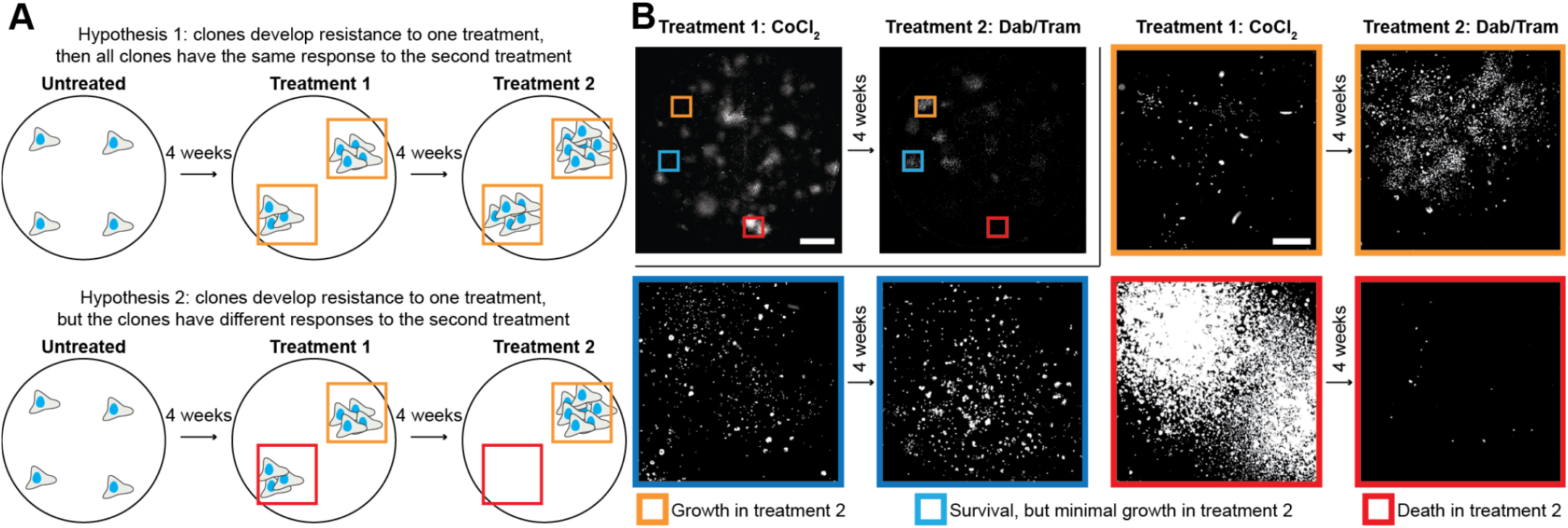
Melanoma cell clones exhibit variable responses to a first and second treatment. **A)** Shown are two possible scenarios for how clones could respond to sequential treatment with different agents. All resistant clones derived from one treatment could have the same response to a second treatment (top), or they could have different responses to a second treatment (bottom). **B)** WM989 BRAF V600E mutant melanoma cells with a nuclear GFP tag imaged after four weeks in CoCl_2_ followed by four weeks in dabrafenib and trametinib (Dab/Tram, top left). The top left images show whole well scans of the clones. The colored boxes show zoomed images selected from the whole well. Orange shows a clone that grew in the second treatment, blue shows a clone that remained a similar size in the second treatment, and red shows a clone that died in the second treatment. Scale bars in whole well scans are 5 mm while scale bars in cropped scans are 500 μm.

To address this gap, we developed an experimental and computational pipeline for long-term tracking of cancer cell clones through sequential and prolonged treatments. We employed DNA barcoding in melanoma cells and used both genomic DNA and single-cell RNA-sequencing (scRNA-seq) to monitor transcriptional and clonal changes over time. We applied a panel of three treatments, allowed four weeks for the development of resistant clones, and then divided samples so that cells from each resistant clone were separately treated again with each of the three treatments. This approach allowed for the same clones to be profiled through all combinations of treatments and treatment orders. Applying this technique, we found that clones exhibit divergent responses to the second treatment. Furthermore, cells within a resistant clone maintain their transcriptional similarity throughout both prolonged treatment with a single agent and sequential treatment with multiple agents. Finally, we demonstrated that gene expression differences between clones underlie a clone’s ability to survive sequential and prolonged treatment.

## Results

### Melanoma cell clones exhibit variable responses to a first and second treatment

To study cancer cell responses through sequential treatments, we selected three conditions relevant to melanoma cells that have diverse mechanisms of action: 1) combined treatment with dabrafenib^19^ and trametinib^20^, which are targeted inhibitors of V600E mutant BRAF and MEK, respectively, 2) CoCl_2_, which mimics hypoxia by activating many of the same stress responses^21,22^, and 3) the chemotherapeutic cisplatin, which kills cells by cross-linking DNA and inhibiting DNA replication^23^. We first investigated whether all clones that develop resistance to one of these treatments exhibit the same response to a second treatment. To visualize individual clones, we sparsely plated BRAF V600E mutant WM989 melanoma cells with a nuclear GFP tag and treated with either dabrafenib and trametinib in combination, CoCl_2_, or cisplatin for four weeks. These treatments resulted in resistant clones that were easily resolved by imaging (Fig. 1B, Supp. Fig. 1). We then applied a different treatment from our panel to these cells for another four weeks. We found that the individual resistant clones originating from the first treatment exhibited diverse responses to the second treatment, consistent with our second hypothesis in Figure 1A. By examining all the clones on the plate, we found that individual clones either grew, survived with minimal growth, or died in a second treatment (Fig. 1B, Supp. Fig. 1). Thus, we concluded that resistant clones originating from one treatment could display highly variable responses to a second treatment. This finding motivated our study of resistant clones at the single-cell level to understand why some survive while others die with sequential treatments.

To build on our initial observation that resistant clones showed variable responses to a second treatment, we sought to generalize this result across a large number of clones and treatment conditions. We used cellular barcoding to allow us to track resistant clones across multiple treatments and through time^9,11,24,25^. We began by uniquely labeling individual cancer cells with a barcoding library developed by Emert et al.^26^ (Fig. 2A). We then allowed cells to undergo a limited number of divisions, producing numerous cells with the same barcode while preserving their gene expression similarities^11,26^. We performed barcode sequencing on a subset of cells to identify clones present before treatment and then divided the remaining population of cells three ways for separate initial treatments with either combination dabrafenib and trametinib, CoCl_2_, or cisplatin. After four weeks of treatment, we performed scRNA-seq and barcode sequencing on a subset of surviving cells and replated the remaining cells from each treatment condition to receive a second round of treatment, with either the same or a different agent, continuing for four more weeks or until reaching confluency. We then repeated scRNA-seq and barcode sequencing of the surviving cells from the second round of treatment (Fig. 2A). At each time point, we also sequenced the barcodes from the genomic DNA (gDNA) of a subset of cells to avoid undersampling the population-level clone dynamics by only including cells captured in the scRNA-seq (see Methods). In summary, this experimental framework yielded quantitative tracking of the clones present before treatment, after the first treatment, and after the second treatment, as well as single-cell and clonally resolved transcriptomics of the resistant cells after the first and second treatments.

**Figure 2:**
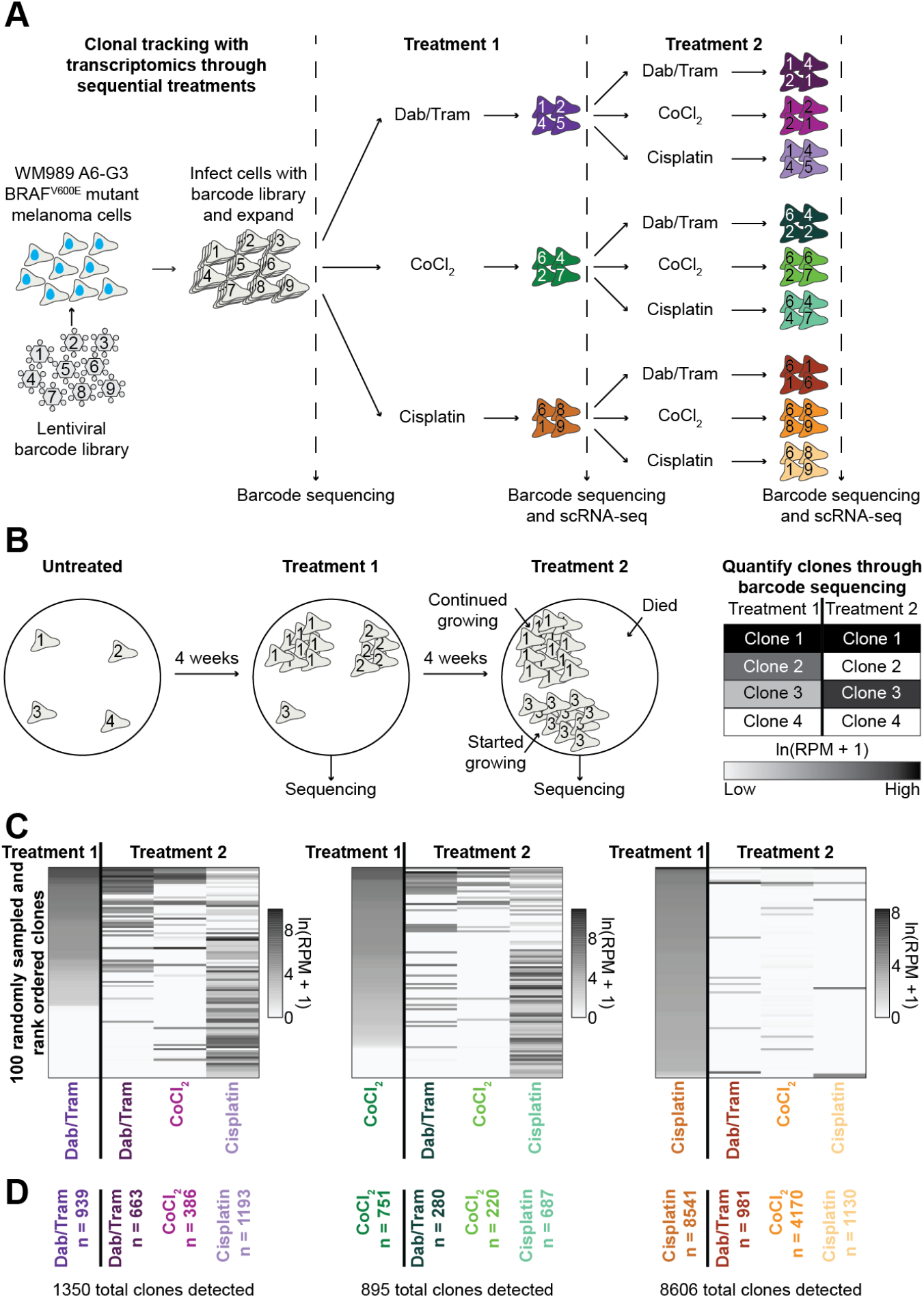
DNA barcoding allows for tracking the transcriptome of clones through multiple rounds of treatment. **A)** Schematic showing experimental protocol for clone tracking and scRNA-seq of WM989 BRAF V600E mutant melanoma cells through treatment with with dabrafenib and trametinib (Dab/Tram), CoCl_2_, or cisplatin (see Methods). **B)** Schematic showing how sequencing reads from the genomic DNA of cells that survived a treatment provide a quantitative readout of treatment response. **C)** Heatmaps showing 100 randomly subsampled and rank-ordered clones detected after initial treatment with Dab/Tram (left), CoCl_2_ (middle), and cisplatin (right) followed by their abundance in the second round of treatment. Individual clones are colored by the ln(Reads Per Million (RPM) + 1) using data from sequencing clonal barcodes from gDNA of surviving cells. **D)** The number of total detected clones per condition are displayed in their assigned color. It should be noted that there are clones whose abundance were below the threshold of detection after treatment 1 that were later detected in treatment 2. The total number of unique clones that were detected after the first or second treatment are displayed below each heatmap.

We first analyzed the gDNA barcode sequencing data to track clones before treatment and after each round of treatment. To equate our sequencing read data to a relative number of cells and set thresholds for resistant clones, we added a “ladder” of barcodes to each sample, consisting of defined numbers of cells with known barcodes (see Methods). Using this ladder, we confirmed that sequencing reads increase with greater cell numbers (Supp. Fig. 2A). Consequently, clones that are highly represented in the population will have a large number of sequencing reads, and those that are lowly represented will have few or no reads (Fig. 2B). We next asked whether clonal growth differences before treatment could explain growth through the first treatment condition (Supp. Fig. 2B). We found that some of the largest clones in the untreated samples were also the largest clones after treatment with each agent. However for each treatment, we also observed clones with low prevalence in the untreated sample, but high prevalence after the first round of treatment. Thus, initial growth differences in untreated clones are not sufficient to explain treatment outcomes.

When we compared the abundance of individual clones between the first and second treatments, we observed three distinct phenomena, which are schematically depicted in Figure 2B. First, we observed clones where the cells survived the first treatment while exhibiting minimal growth, but then grew into large clones during the second treatment. Second, we observed clones that grew significantly during both the first and second treatment. Third, we also observed clones that grew well in the first treatment but then died and were absent after the second treatment. These results reinforce the notion that clonal differences in cells play an important role in response to a second treatment.

When comparing the clones captured after each of the first treatments, we detected a significantly larger number of clones following cisplatin treatment than after combination treatment with dabrafenib and trametinib or CoCl_2_ (Fig. 2C-D). Furthermore, numerous clones with low abundance after initial treatment with dabrafenib and trametinib or CoCl_2_ expanded when cisplatin was applied as the second treatment (Fig. 2C). Although not entirely unique, secondary treatment with cisplatin selected for different clones than secondary treatment with dabrafenib and trametinib or with CoCl_2_. Broadly, these findings highlight the diverse clonal dynamics that can emerge with a single treatment and upon sequential treatments.

### Treatment histories are remembered in gene expression states

Recent studies demonstrate that duration and dose of treatment can influence the transcriptional state of cancer cells^8,9,27^. Therefore, we asked whether the treatment history of multiple agents can be observed in the gene expression of surviving cells, and whether the first or second treatment has a greater effect on the final transcriptional state of a cell. We generated a UMAP projection of the final state of every cell that received two rounds of treatment (hypothetical data in Fig. 3A, actual data Fig. 3B). For each of the three treatments used, we then calculated the pairwise Pearson correlations in gene expression between cells that received the treatment first, and compared them to the Pearson correlations of cells that received that treatment second (Fig. 3B). To provide a baseline for comparison, we randomly sampled a group of the same number of cells from each condition and calculated a Pearson correlation among the random sample. Across every treatment, whether grouped by either the first or second therapy, the average pairwise Pearson correlation of gene expression similarity was at least ~3.5-fold higher than randomly grouped cells. We thus concluded that the final gene expression states of cells that received multiple treatments are influenced by both the first and second treatments. This result was striking, given that these cells had been treated with a second agent, but still retained memory of the first treatment.

**Figure 3:**
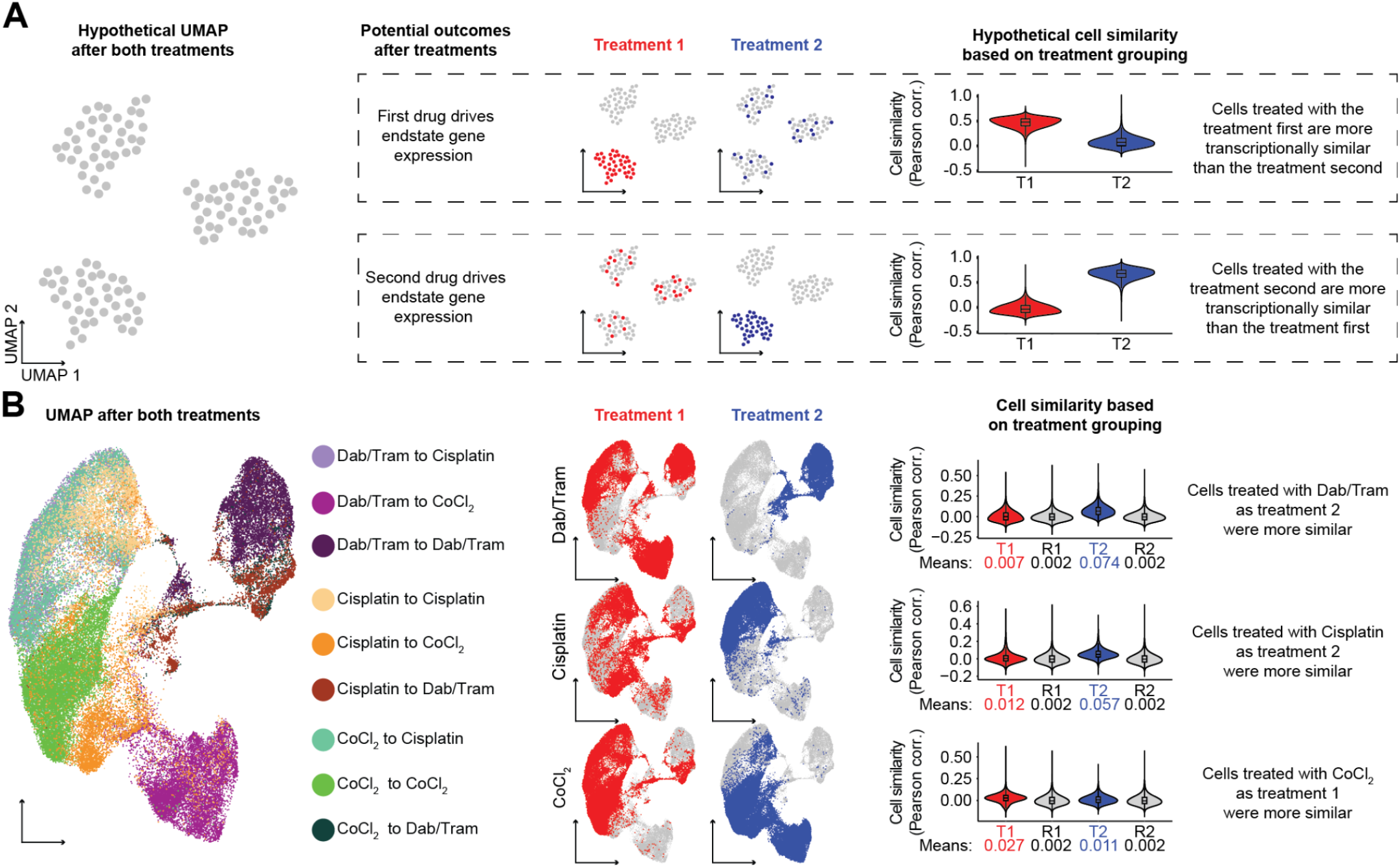
The treatment history of cells are reflected in their gene expression. **A)** Schematic showing the theoretical possibilities of the first treatment driving endstate gene expression (top) or the second treatment driving endstate gene expression (bottom). Displayed are how these possibilities would look in UMAP space or plot as pairwise Pearson correlation between cell transcriptomes. **B)** (Left) UMAP of the nine conditions that had received both a first and second treatment. (Center) UMAP with cells that received the indicated treatment first highlighted in red and those that received the treatment second highlighted in blue. (Right) Quantification of cell similarity using pairwise Pearson correlations of gene expression. For combination dabrafenib and trametinib treatment (Dab/Tram), pairwise Pearson correlations between cells that received Dab/Tram first (treatment 1, T1) were ~3.5-fold higher than the matched random sampling of cells (random control 1, R1), while correlations between cells that received Dab/Tram second (treatment 2, T2) were ~37-fold higher than the matched control (random control 2, R2) and ~10.5-fold higher than those that received Dab/Tram first. For cisplatin treatment, pairwise Pearson correlations between cells that received cisplatin first were ~6-fold higher than the matched random sampling of cells, while correlations between cells that received cisplatin second were ~28.5-fold higher than the matched control and ~5-fold higher than those that received cisplatin first. For CoCl_2_ treatment, pairwise Pearson correlations between cells that received CoCl_2_ first were ~13.5-fold higher than the matched random sampling of cells and ~2.5 fold higher than those that received CoCl_2_ second, while correlations between cells that received CoCl_2_ second was ~5.5-fold greater than the matched random control. All comparisons were statistically significant by a two-sided t-test with p < 2.2e-16. Violin plots display 10,000,000 subsampled data points per condition, but statistical comparisons and averages were calculated on non-subsampled data. Mean values are displayed below each graph.

We next compared the magnitude of the effects of the first and second treatments. Interestingly, we found that secondary treatment with dabrafenib and trametinib (~10-fold greater in average correlation) and Cisplatin (~5-fold greater in average correlation) had a greater effect than initial treatment on gene expression as measured by the Pearson correlation coefficient (Fig. 3B). However, we found the opposite result for cells treated with CoCl_2_ (~2.5-fold less in average correlation). Cells receiving CoCl_2_ as their first treatment were transcriptionally more similar than cells treated with CoCl_2_ as a second treatment. These results collectively indicate that both the initial and subsequent treatments contribute to a cell’s ultimate gene expression state, with the degree of influence varying depending on the treatment.

### Despite treatment-induced changes in gene expression, cells within a clone remain transcriptionally similar to one another

Given that both the first and second treatments influence a cell’s final gene expression state, we next wondered whether these treatment-induced changes effectively eliminated the intraclonal similarities in gene expression that have been previously described in multiple studies^9,11,26^. Thus, we analyzed the cells within each clone to assess if they retained transcriptional similarity after each treatment. For each sample, we identified resistant clones and calculated pairwise Pearson correlations between all cells within each clone (Fig. 4A). We compared these pairwise correlations within clones to correlations calculated on 100 size-matched random samples of cells in the same treatment dataset (Supp. Fig. 3A-L). For every condition, including those from the first treatment and the second treatment, the Pearson correlations within the clone were higher than the randomly sampled data (Fig. 4B-M). As expected, the correlations after prolonged treatment are predominantly lower than their associated four-week sample. However, the pairwise Pearson correlations after the second treatment are still higher than those of randomly sampled cells. Consequently, we concluded that cells within a clone remain transcriptionally similar to each other through sequential and prolonged treatment.

**Figure 4:**
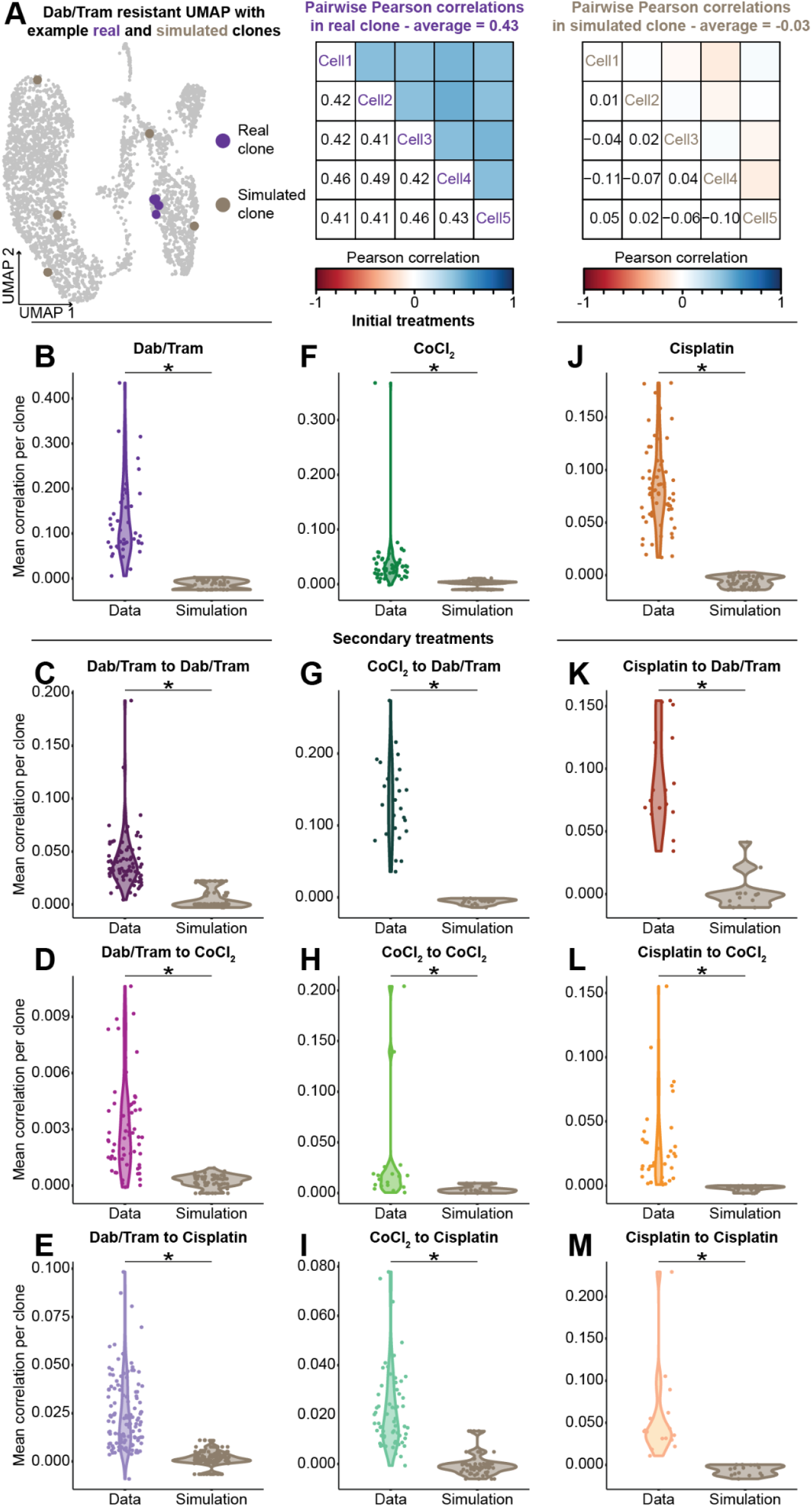
Cells within a clone maintain gene expression similarities through multiple rounds of treatment. **A)** (Left) Location of the cells from a real clone (purple) and its matched random control (tan) from one simulation overlaid on the dabrafenib and trametinib (Dab/Tram) resistant UMAP. (Middle) Pairwise Pearson correlations of gene expression of cells from the real clone. The average of these pairwise Pearson correlations was 0.43. (Right) Pairwise Pearson correlations of gene expression of cells from the simulated clone. The average of these pairwise Pearson correlations was −0.03. For the pairwise Pearson correlation figures, the blocks above the diagonal are shaded to indicate the correlation while the numerical value is displayed in the associated block in the lower diagonal. The averages of the pairwise Pearson correlations of the cells within each clone that received a treatment were compared to those of random samplings of the same number and sized clones over 100 simulations. Clones analyzed ranged from 5 to 11,257 cells. Displayed are the first simulations for each condition. As follows are the number of simulations where the observed similarities were higher than random for each condition by a one-sided Wilcoxon Rank Sum Test with p < 0.05: **B)** Dab/Tram 100/100, **C)** Dab/Tram to Dab/Tram 100/100, **D)** Dab/Tram to CoCl_2_ 100/100, **E)** Dab/Tram to cisplatin 100/100, **F)** CoCl_2_ 99/100, **G)** CoCl_2_ to Dab/Tram 100/100, **H)** CoCl_2_ to CoCl_2_ 100/100, **I)** CoCl_2_ to cisplatin 100/100, **J)** cisplatin 100/100, **K)** cisplatin to Dab/Tram 100/100, **L)** cisplatin to CoCl_2_ 100/100, **M)** cisplatin to cisplatin 100/100.

### Some clones develop induced resistance with sequential treatment

Our experimental design, which started with a large pool of clones to test many sequential treatment combinations, provided a unique opportunity to identify clones that only survived a particular treatment after developing resistance to a different treatment beforehand – a phenomenon we termed “induced resistance” (Supp. Fig. 4A). In each ordered combination of treatments, we identified hundreds of clones with induced resistance (Supp. Fig. 4B,G,H). To measure the gene expression differences in these induced resistant clones, we performed differential gene expression and gene set enrichment analysis comparing clones with induced resistance to clones that were sensitive to the second treatment (Supp. Fig. 4B,C,G,H,I,J). We discovered significant transcriptional and pathway level differences in the induced resistant clones (Supp. Fig. 4C,I,J). One such pathway enriched in clones that only survived CoCl_2_ after developing resistance to dabrafenib and trametinib was IL6/JAK/STAT3 signaling (Supp. Fig. 4C), of which IL6ST is an upstream receptor whose gene is also highly enriched^28^ (Supp. Fig. 4D,E). IL6, which signals through IL6ST, has previously been implicated in chemotherapy and targeted inhibitor resistance in multiple cancer types^29–32^. We observed that *IL6ST* is infrequently expressed prior to treatment (expressed in 2.1% of cells, Supp. Fig. 4F), but more frequently expressed after survival in dabrafenib and trametinib (expressed in 44.2% of the cells, Supp. Fig. 4E). These findings suggest that treatment with dabrafenib and trametinib can cause previously CoCl_2_-sensitive clones to become resistant, with this resistant state marked by expression of *IL6ST*.

### Prolonged treatment with the same agent causes population-level and clonal changes in gene expression

We next investigated the gene expression changes in cells undergoing prolonged treatment with a single agent. We found that cells clustered based on the duration of treatment across our entire panel (Fig. 5A). To assess pathway-level changes, we performed differential gene expression comparing samples at the first and second time points for each treatment. We then used enrichGO33 to identify enriched pathways and grouped GO terms that were related to similar cellular functions (Fig. 5B, Supp. Fig. 5A-H). In all three treatments, we found that prolonged treatment resulted in significant changes in pathway activity, most of which have been reported in the literature to be involved with treatment resistance or aggressive cancer phenotypes^34–41^. Thus, across all three different treatments, we found further evidence that the transcriptional states of the population change dynamically as a function of the duration of treatment^27^.

**Figure 5:**
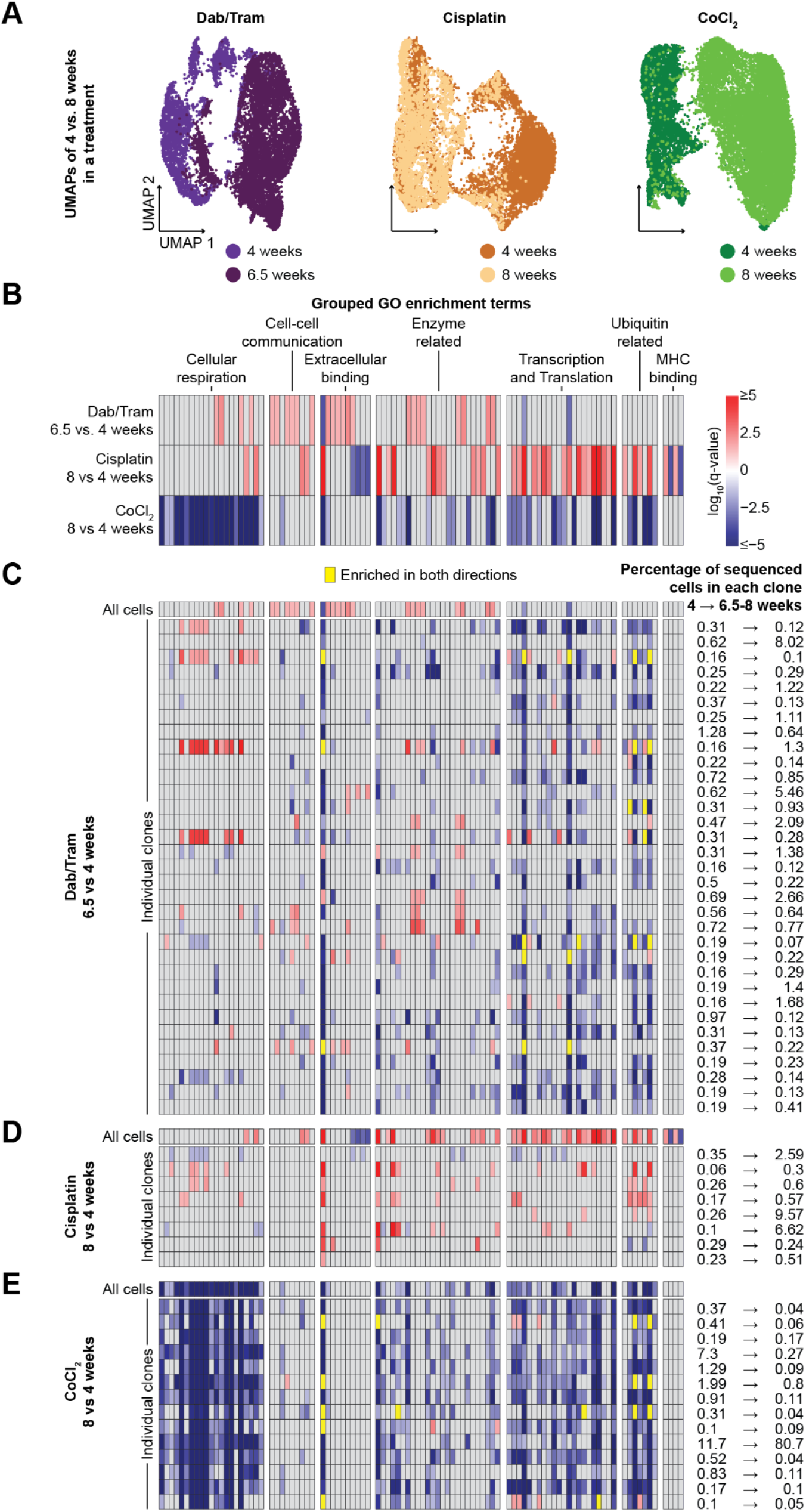
Prolonged treatment with the same agent causes population-level and clonal changes in pathway expression. **A)** UMAPs comparing cells sequenced after four and 6.5-8 weeks in treatment with (left) dabrafenib and trametinib (Dab/Tram), (middle) cisplatin, or (right) CoCl_2_. **B)** We identified differentially expressed genes between 6.5-8 and four weeks of treatment with each agent. We then identified GO terms enriched after either the first round of treatment (blue) or after the second round of treatment (red) based on the log_10_(q-value) and -log_10_(q-value) respectively. GO terms that were not significantly enriched, or failed other thresholds, in either direction for a condition are colored in gray. Enriched pathways were grouped into categories of related function, and the pathways that make up each group are detailed in Supplemental Figure 4. Subpanels **C-E** perform perform the same analysis as performed in **B** with all of the cells from each condition (with that comparison displayed again) on a clone-by-clone basis for all clones with at least five cells in both rounds of treatment for Dab/Tram, cisplatin and CoCl_2_, respectively. On the right of each subpanel is displayed the percentage of total sequenced single-cells per condition from each clone after each round of treatment.

Considering that treatment duration produced differences in transcriptional states, we identified two possible explanations for this observation. One possibility is that all clones undergo the same population-level changes in gene expression during treatment. Alternatively, individual clones might have heterogeneous changes during treatment, which collectively contribute to the average signal observed at the population-level. To explore these possibilities, we performed the same pathway analysis at the clonal level (Fig. 5C-E). In targeted therapy with dabrafenib and trametinib, the majority of pathways that were differentially active between four and 6.5 weeks of treatment at the population-level were not consistently differentially expressed on a clone-by-clone basis (Fig. 5C). Similarly, pathways that were differentially active between cisplatin treated cells after four and eight weeks at the population-level were not reflected in most of the individual clones (Fig. 5D), suggesting that clones can have differential responses to prolonged treatment with the same agent.

While the population-level changes with combination dabrafenib and trametinib or cisplatin were largely not evident in the majority of clones, we found that CoCl_2_ showed the opposite result. Almost all CoCl_2_ clones displayed the same pathway enrichment as the analysis performed at the population level (Fig. 5E). Interestingly, despite all of the clones going through similar gene and pathway expression changes, we still observed drastic changes in cell growth between clones, with one clone constituting 81% of the total cells sequenced after eight weeks in CoCl_2_. Across all three treatments, the one pathway where the population-level changes were reflected in all of the clones was cadherin binding, yet the directionality of this change varied across treatments. These findings demonstrate that different treatments can elicit different responses from individual clones.

### Clonal differences in gene expression underlie survival through prolonged treatment with dabrafenib and trametinib

Given that individual clones can have divergent responses to treatment, we next wondered whether the specific gene expression state of clones at the first point could predict which clones will be resistant at a later time. We focused specifically on combination treatment with dabrafenib and trametinib. Previous work from our lab showed that *EGFR* and *NGFR* are both marker genes which are highly predictive of survival to dabrafenib and trametinib at four weeks^11^. Our scRNA-seq data of dabrafenib and trametinib resistant cells showed two independent clusters of resistant cells, one which was high in *EGFR* and another which was high in *NGFR* (Supp. Fig. 6A-D). We hypothesized that these divergent transcriptional states correlated with differences in survival to prolonged treatment with dabrafenib and trametinib. After four weeks of treatment with dabrafenib and trametinib, we identified 44 resistant clones, 23 of which were exclusively in the *EGFR*-high cluster, five of which were exclusively in the *NGFR*-high cluster, and 16 of which had cells in both clusters (Fig. 6A). We then followed these 44 clones through an additional 2.5 weeks in dabrafenib and trametinib. We found that 13.0% and 18.8% of the *EGFR*-high and mixed clones died, respectively, (as defined by no longer passing resistant thresholds, see Methods) while 100% of the *NGFR*-high clones died (Fig. 6A), suggesting that *NGFR*-high clones are able to survive targeted therapy on short, yet not on prolonged, timescales. Interestingly, when looking at these same clones after treatment with dabrafenib and trametinib followed by CoCl_2_, many of the *NGFR*-high clones also died (Supp. Fig. 6E-F). Conversely, the majority of these 44 clones that had any growth after treatment with dabrafenib and trametinib followed by cisplatin were originally *NGFR*-high or mixed, indicating that *NGFR* enrichment may indeed be beneficial for subsequent survival through cisplatin (Supp. Fig. 6G). The enrichment and depletion of clones with specific expression patterns further supports the idea that gene expression states confer treatment-specific survival consequences

**Figure 6:**
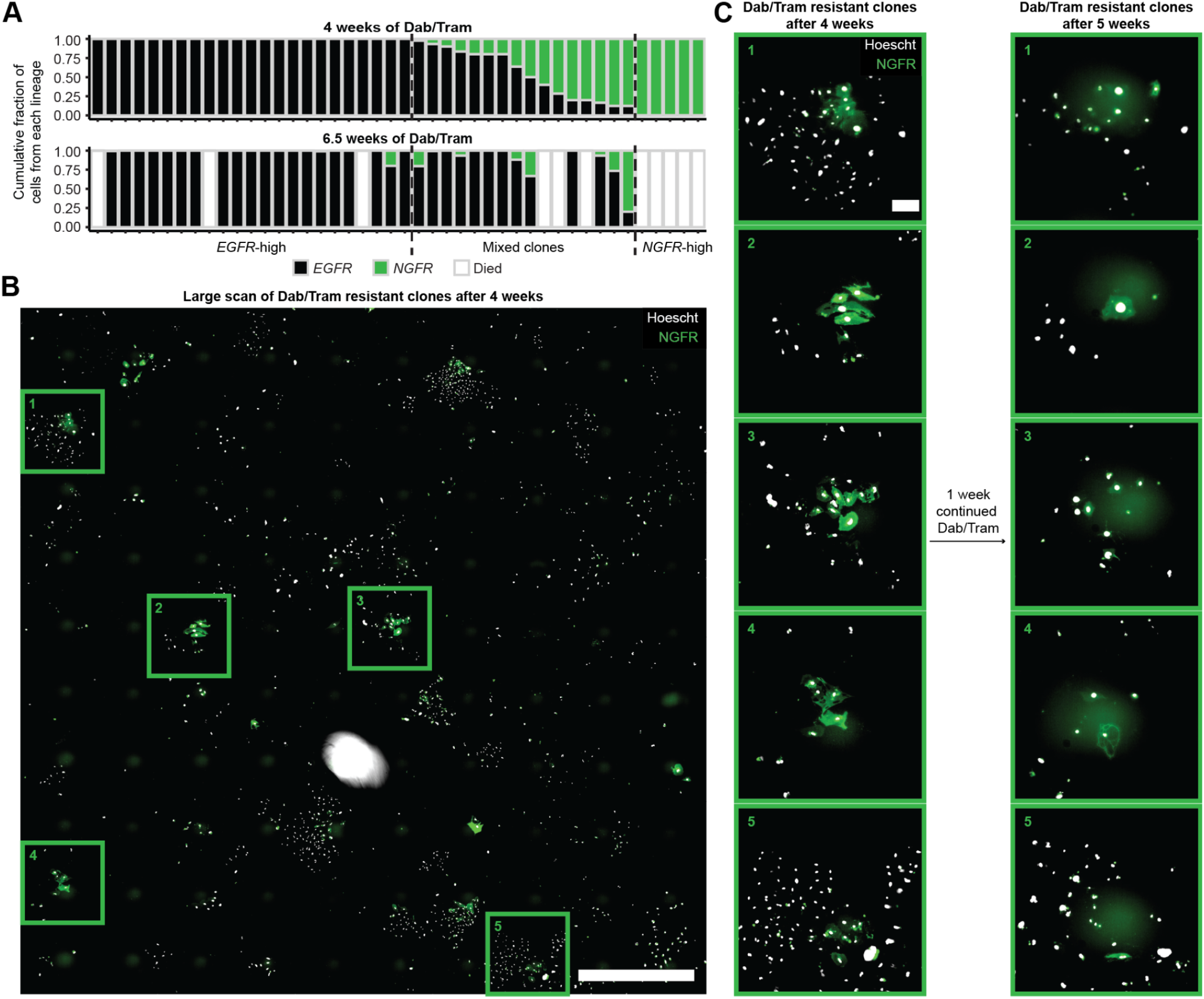
Clonal differences in gene expression underlie survival to prolonged treatment with dabrafenib and trametinib. **A)** Stacked colored bars represent the proportion of cells from each clone with at least five cells after initial treatment (four weeks) with dabrafenib and trametinib (Dab/Tram) in the (black) *EGFR*-high cluster or (green) *NGFR*-high cluster (top). Dotted lines separate clones that are all *EGFR*-high, mixed *EGFR* and *NGFR* expression, and *NGFR*-high. Aligned below are the same clones after the second round of treatment (6.5 weeks) with Dab/Tram. Clones that still have greater than five cells are again colored by their proportion of cells in either the *EGFR-* or NGFR-high cluster while clones that have died (less than five cells) are colored white. **B)** Large scan of NGFR-mNG2(11) WM989 A6-G3 cells imaged after initial treatment with Dab/Tram. Scale bar is 2 mm. **C)** Cropped scans highlighting five NGFR-mNG2(11) WM989 A6-G3 clones imaged after initial treatment with Dab/Tram and again one week later. Positions of cropped scans are boxed and numbered in subpanel **B**. All cropped scans of the same time point are contrast matched. The two timepoints were not contrast matched, and rather were contrasted separately to visualize faint signal. Scale bars are 200 μm. For subpanels **B** and **C**, Hoechst stained nuclei are displayed in gray and NGFR-mNG2(11) is displayed in green.

To validate our finding that *NGFR*-high clones died in prolonged treatment with dabrafenib and trametinib, we used WM989 melanoma cells with NGFR tagged at the endogenous locus with an mNeonGreen2^42^. We treated these tagged cells with dabrafenib and trametinib for four weeks and saw the emergence of green (NGFR-high) colonies and colonies that were not expressing NGFR (presumably EGFR-high) (Fig. 6B), confirming the presence of colonies with NGFR expression. At five weeks in dabrafenib and trametinib, we found that many of the colonies that previously contained NGFR-expressing cells had died, while adjacent cells with lower NGFR expression survived (Fig. 6C). This result further supports that high *EGFR* expression is more indicative of long-term resistance to dabrafenib and trametinib, while high NGFR expression is associated with shorter-term survival. More broadly, these data demonstrate that clonal differences in gene expression can predict clonal outcomes to prolonged treatment.

## Discussion

Tumor heterogeneity has been extensively studied for the role it plays in treatment resistance^1,2,5–11^; however, most of these studies have focused on resistance to a single treatment^2,4–8,10^. Sequential treatment is a common approach for cancer treatment, but has predominantly been studied through clinical trial data or at the bulk tumor/cell population level1^6,43–48^. Here, we focus on the single-cell and single-clone level, addressing how resistant clones change in their abundance and transcriptional state over the course of sequential treatments. We combine imaging, scRNA-seq, and clonal tracking with cell barcoding to discover that individual resistant clones show highly variable responses to a second treatment, which highlights the importance of studying sequential treatment resistance at a clonal, rather than population level.

Our findings further support a growing body of evidence that resistance to treatment can not be explained by single mechanisms acting in isolation^10,49^. Rather, resistance reflects a complex and dynamical process in which diverse and continuously evolving cellular states are generated^9,27^. Our work contributes another layer of phenotypic complexity by demonstrating that the diverse clones generated under treatment have differential sensitivities to subsequent treatments. We further connect the response of individual clones to specific gene expression states, thus supporting the notion that response to sequential treatments is nonrandom and occurs at the clonal level. The connection between phenotype and gene expression is similar to observations before treatment, where preexisting differences in the initial state of cells are predictive of treatment outcome^8,26^.

This study also contributes to our understanding of gene expression memory. While individual resistant clones have a diversity of transcriptional states, the cells within these clones are transcriptionally similar to each other. These states are heritable, such that the cells derived from a common parent cell have related transcriptional profiles. Such heritability has been measured in the absence of treatment and upon a single agent^9,11,26,42^, but here heritability of gene expression extends through prolonged and sequential treatments. We speculate that these clonal gene expression similarities are epigenetically encoded, but the full mechanisms remain to be elucidated. Furthermore, this heritability of gene expression is critical for our experimental design as it enables comparisons between the same clones receiving different treatments. Given that clones are still transcriptionally similar after the second treatment, this experimental and analytical framework could be easily scaled to include third and potentially even fourth rounds of treatment for assessing clonal dynamics on longer timescales and with additional treatments.

While our experimental design captures large resistant clones after treatment, it is less sensitive for smaller clones as our data is highly subsampled. Importantly, the experimental framework relies on the ability of clones to proliferate for them to be captured at the different time points and evenly distributed across the samples. Thus, this data is not able to comment on non-proliferative states and/or very slow cycling states, which have been implicated in pre-existing states associated with resistance^4,5,50,51^. While clones consisting of only one cell can be detected in our data and others^9^, they are not likely to be present across multiple time points or treatment conditions. Rather, because our experimental design emphasizes the most proliferative clones, we are best equipped to define how large clones respond and change during sequential treatment. Subsequent studies and alternative experimental designs would be needed to better capture the role of these small clones in resistance.

In summary, we find that intraclonal similarities in gene expression are preserved over multiple months and across different treatments. Moreover, we show that interclonal differences in gene expression inform survival through sequential and prolonged treatment, suggesting that the progression of resistance in these treatments is nonrandom. This knowledge lays the foundation for future work to target clonal states to prevent resistance to sequential and prolonged treatment.

## Methods

### Treatments

We made stock solutions of 500 μM dabrafenib (Cayman, 16989-10), 5 μM trametinib (Cayman, 16292-50 and SelleckChem, S2673 871700-17-3), and 25 mM CoCl_2_ (Spectrum Chemical Manufacturing Corporation, 18609836) as follows: dabrafenib, trametinib in DMSO, and cisplatin in 154 mM NaCl, CoCl_2_ in nuclease-free water. For the single-cell experiments, we diluted all agents in culture medium to a final concentration of 15 μM cisplatin, 200 μM CoCl_2_, and 2.5 nM trametinib plus 250 nM dabrafenib for combined dabrafenib/trametinib treatment.

### Cell lines and tissue culture

We performed all experiments on WM989 A6-G3 melanoma cell lines, including variations with a GFP nuclear reporter (H2B-GFP) and a split mNeonGreen2 NGFR tag (NGFR-mNG2(11))^42^. WM989 A6-G3 cells were derived from WM989 cells and single-cell bottlenecked twice. WM989 A6-G3 cell lines were authenticated and validated as mycoplasma negative^11^. We cultured all WM989-derived lines in TU2% media consisting of 78% MCDB 153, 20% Leibovitz’s L-15, 2% FBS, 1.68 mM CaCl_2_, 50 U/mL penicillin, and 50 ug/mL streptomycin. We passaged non-treated cells with 0.05% trypsin-EDTA, and treatment-resistant cells were passaged with 0.25% trypsin-EDTA. For lentivirus packaging, HEK293T cells were cultured in DMEM containing 10% FBS, 50 U/mL penicillin, and 50 ug/mL streptomycin. We passaged HEK293Ts with 0.05% trypsin-EDTA. All cells were maintained at 37°C and 5% CO_2_.

### Treatment resistant colony experiments

We treated H2B-GFP WM989 cells with one of each of the following treatments for four weeks. For cells treated with 200 μM CoCl_2_ in Tu2% medium, we performed a media change every three to four days, each time treating with CoCl_2_ treated media. For cells treated with 15 μM cisplatin, we treated with cisplatin treated TU2% medium for the first three days, then every following media change used normal TU2%. For cells treated with 250 nM dabrafenib and 2.5 μM trametinib, we performed a media change every three to four days, each time treating with dabrafenib and trametinib treated TU2% medium. For the next four weeks of treatment, we treated cells with a second round of treatment with cisplatin, CoCl_2_ or a combination of dabrafenib and trametinib using the corresponding protocol for weeks five through eight. We imaged cells at four weeks (following the first round of treatment) and eight weeks (following the second round treatment) in the GFP channel (20 ms exposure). Images were contrasted using custom python scripts.

### Barcode Library

We performed DNA barcoding with the Rewind library using barcode plasmids provided generously by the Raj Lab at the University of Pennsylvania and produced as previously described^26^. Briefly, PAGE-purified Ultramer oligonucleotides were ordered containing a 100-base pair sequence repeating ‘WSN’, where W = A or T, S = G or C, and N = any base pair, and inserted into a lentivirus vector backbone downstream of the EFS promoter and GFP sequence. Emert et al. have published a detailed protocol of how these barcodes are produced (https://www.protocols.io/view/barcode-plasmid-library-cloning-5qpvon6yxl4o/v1)^26^, and the plasmid sequence is available (https://benchling.com/shaffer_lab/f/lib_6JXLhfQH-plasmids/seq_BUIxqCk0-lentiefs_gfp_100bp_barcode_v1/e_dit?m=slm-GJ609ijArVWmkT8mk8zr)^11^.

### Lentiviral packaging

We packaged barcodes into lentivirus following the procedure outlined for the Rewind library^26^. In brief, HEK293T cells were cultured in three 10 cm plates until nearing confluency. For all three plates plus one half a plate worth of excess solution, we combined 1750 μL Opti-MEM (Gibco, 31985062) with 280 μL PEI in one tube while another 1750 μL Opti-MEM was mixed with 35 μg (29.2 μL) barcode plasmid, 26.25 μg (20.9 μL) pPAX2, and 17.5 μg (30.2 μL) VSVG in a second tube. We incubated each tube at room temperature for five minutes before combining them, vortexing, and incubating for an additional 15 minutes. We then slowly added 1106 μl of the mixed solution to each of the three 10 cm plates. After six hours, we discarded the media and replaced it with seven ml of DMEM. After 24 hours and confirming GFP expression, we collected the virus-containing media into a conical tube stored at 4°C and replaced it with another seven mL of fresh media. After collecting virus twice more at 24 hour intervals, we centrifuged the virus-containing media at 3000 rpm for five minutes. We collected the supernatant and passed it through a 0.45 μm filter (Millipore Sigma, SE1M003M00). Finally, we made 1 mL aliquots and placed them at −80°C for storage.

### Lentiviral transduction

For experiments involving the transduction of WM989 A6-G3 with barcode virus, we seeded cells in six-well plates at 300,000 cells per well and dropwise added 32 μL (as determined by titration for 20-25% infection efficiency) of freshly thawed virus. We then centrifuged the cells at 1750 RPM for 25 minutes and incubated overnight at 37°C. The following morning, we removed the media, washed the cells once with DPBS, and added fresh TU2% media.

### Fluorescence-activated cell sorting (FACS)

Three days after transducing cells with the GFP-barcode virus, we performed fluorescence-activated cell sorting to isolate GFP-expressing, barcoded WM989 A6-G3 cells. We used 0.05% trypsin-EDTA to obtain single-cell suspensions, neutralized with TU2%, washed the cell pellet with 1% BSA-DPBS, and resuspended in 1% BSA-DPBS containing DAPI. We passed the resuspended cells through a FACS cell strainer (Falcon, 352235) prior to FACS. We performed sorting on a Beckman Coulter MoFlo Astrios sorter with a 100 μm nozzle. We gated for singlets and live cells, then sorted for GFP-positive cells.

### Single-cell RNA sequencing experiment

For the single-cell experiment outlined in Figure 2A, we trypsinized cells that had been sorted for being GFP-barcode positive the previous day and replated 425,000 starting cells. We allowed these cells to double roughly 5.5 times so that we had ~45 cells per clone. We pelleted and froze ⅓ of these cells at −80°C for later barcode sequencing from gDNA. We then plate 225,000 cells in 18 x 10 cm plates each (54 total) for treatment with either a combination of dabrafenib and trametinib, CoCl_2_, or cisplatin,. The following day, we began treating the cells. For combination treatment with dabrafenib and trametinib and for treatment with CoCl_2_, treated media was added every three to four days for four weeks. For cisplatin treatment, we treated cells for 72 hours before replacing with normal media which was maintained until the end of four weeks. After four weeks, resistant cells were trypsinized. From each treated population, we removed ¼ of the surviving cells for scRNA-seq of ~7,000 cells and barcode sequencing from gDNA of the remaining cells. The remaining cells from each population were split evenly to receive the three secondary treatments for an additional four weeks, as previously described. After the second round of treatment was complete, we performed scRNA-seq on ~10,000 cells and froze the remaining at −80°C for barcode sequencing from gDNA. One sample, combination treatment of dabrafenib and trametinib into dabrafenib and trametinib, collected after only 2.5 weeks of secondary treatment as the cells were nearing confluency.

We performed all scRNA-seq using the 10X Genomics 3’ v3.1 Dual index Single-Cell gene expression kit on a 10X Chromium Controller. We quantified libraries using the Qubit dsDNA High Sensitivity Assay (Invitrogen, Q32854) and the Agilent Bioanalyzer High Sensitivity DNA Kit (Agilent, 5067-4626). We then diluted libraries to four nM and pooled samples. We sequenced pooled samples on a NextSeq 500 with 75 cycle high output kits (Illumina, 20024906) using 28 cycles for read one, 10 cycles for each index, and 43 cycles for read two.

### DNA barcode recovery from scRNA-seq

We amplified DNA barcodes from excess full-length cDNA generated during the 10X workflow as described previously^11^. In brief, we performed targeted amplification of the barcode from the full-length cDNA using primers with Illumina adapter sequences, sample indices, and staggered bases of different lengths from Harmange et al. and Goyal et al.^9,11^ (Supp. Table 1). For each sample, we diluted cDNA to 8.75 ng/μL and performed PCR reactions using 5 μL cDNA, 500 nM primers, 25 μL NEBNext Q5 HotStart HiFi PCR Master Mix (NEB, M0543S), and nuclease-free water for a total volume of 50 μL. We ran reactions on a thermal cycler using the following settings: 98°C for 30 seconds, 13 cycles of 98°C for 10 seconds and 65°C for two minutes, followed by a final step at 65°C for five minutes. After PCR, we purified the libraries using 0.7x (35 μL) AMPure XP magnetic beads (Beckman Coulter, A63881) followed by two 80% ethanol washes and a final elution of our purified barcodes in 20 μL of nuclease-free water per sample. We quantified purified libraries using Qubit dsDNA High Sensitivity Assay (Invitrogen, Q32854) and the Agilent Bioanalyzer High Sensitivity DNA Kit (Agilent, 5067-4626). We diluted libraries and sequenced an equimolar pool all samples on a NextSeq 500 using a 150 cycle mid output kit (Illumina, 20024904) using 28 cycles for read one to capture the 10X barcodes and UMIs, eight cycles for each index, and 123 cycles on read two to capture the clone DNA barcodes.

### Creating DNA barcode ladders

To create standardized ladders to relate gDNA barcode reads to number of cells, we used FACS as described above to sort barcoded, GFP-positive single cells into each well of four 96-well plates and cultured the cells in untreated TU2% media as described above. These single cells were allowed to expand into single-clone colonies which we grew to confluence in 10 cm plates, at which point we harvested a portion of each clone for barcode sequencing. We left the remainder to expand further. We performed barcode library preparations on the harvested cells from each clone as described below by isolating gDNA, PCR amplifying the barcode sequence, and performing double-sided bead cleanup. These sequences were then sent for Sanger sequencing (Eurofins Genomics). From the ten clones sequenced, we selected six to use as ladder clones which contained correct conserved regions and had no ambiguous bases. We selected two clones each to be high, medium, and low abundance clones in the ladder, corresponding to 1000, 500, or 50 cells. We then trypsinized and counted live cells from each clone via hemocytometer and added the correct number of cells from each clone into tubes which were stored at −80°C.

### DNA barcode recovery from gDNA

We performed barcode library preparations from genomic DNA (gDNA) as previously described with slight modifications^26^. In brief, we thawed cell pellets that had been stored at −80°C and added a “ladder” of cells with known barcode sequences and known initial cell numbers (two replicates each of 50, 500, and 1000 cells per clone) to each pellet (Supp. Fig. 2A). We then isolated gDNA from cells using the QIAmp DNA Mini Kit (QIAGEN, 51304), following the spin protocol for DNA purification from blood or bodily fluids. We then amplified DNA barcodes from gDNA using primers with Illumina adapter sequences, sample indices, and staggered bases of different lengths from Harmange et al.^11^ (Supp. Table 1). For each sample, we performed several PCR reactions (utilizing 20-40% of the total isolated gDNA), each containing 500 ng of gDNA, 500 nM primers, 25 μL of the NEBNext Q5 HotStart HiFi PCR Master Mix (NEB, M0543S), and nuclease-free water to a final volume of 50 μL. We ran reactions on a thermal cycler using the following settings: 98°C for 30 seconds, 23 cycles of 98°C for 10 seconds and 65°C for 40 seconds, and 65°C for five minutes. After PCR, we pooled individual reactions from the same sample before cleaning. We performed double-sided library purifications on the pooled samples using first 0.6x (30 μL x the number of pooled reactions) AMPure XP magnetic beads (Beckman Coulter, A63881) and keeping the supernatant. To the supernatant, we added 1.2x (30 μL x the number of pooled reactions) AMPure XP magnetic beads and washed twice with 80% ethanol. We eluted our purified barcodes from the beads using nuclease-free water (20 μL x the number of pooled reactions). We quantified purified libraries using Qubit dsDNA High Sensitivity Assay (Invitrogen, Q32854) and the Agilent Bioanalyzer High Sensitivity DNA Kit (Agilent, 5067-4626). We diluted and made an equimolar pool of all libraries for sequencing on a NextSeq 500 using a 150 cycle mid output kit (Illumina, 20024904) using 151 cycles for read one to capture the clone DNA barcodes and eight cycles for each index.

### Clone abundance heatmaps from gDNA

We calculated the ln(Reads Per Million (RPM) + 1) of clones with at least two reads in the pre-treatment sample or with reads greater than one of the two corresponding 50-cell ladder clones in a given initial treatment sample or any subsequent treatment samples. We then subsampled and rank-ordered 100 clones for display in the heatmaps in Figure 2C and Supplementary Figure 2B and displayed the number of clones with at least one sequencing read.

### scRNA-seq analysis

We generated gene count matrices using the CellRanger software 5.0.1 (10X Genomics) with the 2020-A hg38 reference genome, and we performed downstream analyses in the Seurat v4 package^52^. We also used CellRanger Feature Barcode technology to merge our custom clone barcodes sequencing with scRNA-seq data. For each sample, we filtered to remove cells with high RNA count indicating doublets, low numbers of genes detected, and a high percentage of mitochondrial reads. Data was processed using NormalizeData, FindVariableFeatures, ScaleData, RunPCA, FindNeighbors, FindClusters, and RunUMAP in that order. Downstream analyses are described in their associated methods sections.

For calculations of the percentage of cells in a condition expressing a gene, a cutoff of greater than one count was used.

### Combining scRNA-seq and barcode data

To identify clone barcodes present in our sequencing data, we used a custom pipeline described by Harmange et al.^11^ and available here: https://github.com/SydShafferLab/BarcodeAnalysis. Briefly, this pipeline takes in FASTQ files containing sequencing reads of barcodes from both gDNA and cDNA and identifies unique barcode sequences throughout all files. This code then collapses highly similar sequences with a Levenshtein distance of eight or less that are likely the same barcode which contains small alterations caused by small point mutations, PCR artifacts, or sequencing artifacts. The pipeline outputs a reference file along with both gDNA and cDNA FASTQ files that were corrected for collapsed barcodes. The corrected cDNA FASTQ files and the reference file can be input into the CellRanger Feature Barcode pipeline to generate count matrices and link cDNA barcode sequencing reads with their cell of origin based on the 10X barcode and UMI. Additionally, this pipeline separately outputs a list of total counts per barcode from either the cDNA and gDNA samples sequenced from each condition .

After combining scRNA-seq with cDNA barcode sequencing data, we found that some cells had reads from multiple barcodes. To assign the dominant barcode to each cell, we separately set a reads-based cutoff for each sequencing sample that maximized the total number of cells that only had a single barcode above the cutoff number of reads. In the case that a cell contained multiple barcodes each with reads above the cutoff, we assigned that cell a dominant barcode only if the reads of the highest detected barcode were 3-fold higher than the second highest detected barcode. Cells without a dominant barcode were excluded from clone-based analyses.

### Determining resistant clones for single-cell analysis

From the gDNA reads, we identified resistant clones in each sample as any clone with more reads than the lower number of reads associated with one of the two 50-cell ladder clones in that sample. We supplemented these lists by adding lineages which had at least one read in the sample as well as more than 50 cells-equivalency gDNA reads within any sample from the same sequential treatment group (e.g. in either cisplatin, cisplatin to cisplatin, cisplatin to CoCl_2_, or cisplatin to dabrafenib and trametinib for any within this group) to capture the prevalence of clones which expanded in related treatment groups and were likely to be found in the population. We then supplemented the above lists with clones which had more than 50 cells in at least one of the four samples from the corresponding sequential treatment group for each cDNA sample and also had at least one read in the gDNA sample. This ensured that we still analyzed clones that had grown out in one of the related treatment groups, even if it only had minimal gDNA representation.Finally, we subsetted these lists to contain only clones with at least 5 cells in each individual cDNA sample. These methods ensured that all resistant clones had some level of detection in both gDNA and cDNA barcode sequencing without filtering out important clones that were only highly detected in later treatment groups.

### Analysis of transcriptional similarity based on treatment order

We grouped cells from samples that had survived 6.5-8 weeks of any combination of treatment based on whether they received a combination of dabrafenib and trametinib, CoCl_2_, or cisplatin as either the first or second treatment. We then calculated pairwise Pearson correlations based on the scaled data generated in Seurat^52^ of 2000 variable genes for all cells within the group. Additionally, we created random datasets matching the same number of cells for each experimental grouping of cells and calculated the pairwise Pearson correlations identically. We then performed two-tailed t-tests comparing the pairwise Pearson correlations of cells grouped by receiving a treatment as either their first or second treatment. We additionally compared each grouping of cells to the associated random dataset. For plotting violin plots of pairwise Pearson correlations, we subsampled 10,000,000 random data points.

### Clone similarity analysis

We first identified resistant clones with at least five cells in each of the 12 treatment conditions (as described above). We then calculated the pairwise Pearson correlations in gene expression across the scale.data^52^ of 2000 variable genes within the cells from each clone in each condition. Meanwhile, for each clone in each condition, we randomly sampled size-matched controls from cells that had survived the same treatment 100 times. For each simulation, we compared the average Pearson correlations of the real clones to those of the simulated clones using a one-sided Wilcoxon Rank Sum Test to test whether the correlation within the real clones were greater than the simulated clones. In Figure 4, the first simulation for each treatment condition is displayed.

### Induced resistance analysis

To find clones with resistance that was induced to survive a treatment by first surviving a different treatment, we identified resistant clones within each treatment condition with a lenient filter of having more than one sequencing read in the barcodes sequenced from gDNA. The more lenient determination of whether a clone was resistant to a condition ensured that we accurately identified clones that had not survived a treatment from those that had survived a condition with only minimal growth such that it was not previously identified as resistant. We then generated lists of induced clones that only survived a treatment after developing resistance to a different treatment. We then mined these lists for clones for those which we had corresponding scRNA-seq data. We then used FindMarkers in the Seurat package^52^ to identify genes up- or downregulated in induced clones compared to cells from clones that had survived the same initial treatment, but were not induced to survive the second treatment (Supp. Fig. 4B,G,H) with a logfc.threshold of 0.25 and p_val <= 0.05. We then performed directional gene set enrichment of the Molecular Signals Database v2022.1 collection of hallmark gene sets53,54 using the fgsea package (v1.22.0)^55^ with a minSize of 5 and maxSize of 500.

### Gene ontology analysis

We used the FindMarkers method in Seurat^52^ to create two lists of genes that increased or decreased between the four and 6.5-8 week time points in the same treatment. We input universally normalized Seurat objects (see above). We used default parameters for Seurat objects. We only listed genes with a minimum log^2^ fold change greater than 0.25 and that were expressed in at least 10% of cells in one. We added 1 to average expression values when log_2_ fold change was calculated. We input these lists into the enrichGO function in the clusterProfiler^33^ library using default parameters and a p-value cutoff of 0.05. We used Bonferroni corrections to adjust p-values and only analyzed gene ontologies with q-values of 0.2 or less. Further, we only considered gene sets with a minimum of 10 genes and a maximum of 500 genes. We displayed gene ontologies that did not pass thresholds in gray and gene ontologies that went up in both directions as yellow. We determined “molecular function” gene ontologies. We plot GO terms based on genes whose expression increased over time as the -log_10_(q-value) (resulting in positive values) and those based on genes whose expression decreased over time as the log_10_(q-value) (resulting in negative values). We analyzed resistant clones with five or more cells (as described above) at both time points using the same methodology as applied to whole populations to determine GO terms that had changed within individual clones.

### Computational analysis of clone abundance during continued treatment of *EGFR-* and *NGFR*-high dabrafenib and trametinib resistant clones

To assess how prolonged combination treatment with dabrafenib and trametinib affected clones with high *EGFR* or *NGFR* expression, we began by clustering the cells sequenced after four and 6.5 weeks of dabrafenib treatment together and forced Seurat to cluster these cells into only two subclusters using FindClusters with a resolution of 0.015. Based on gene expression plots, we identified one cluster as *EGFR*-high and one cluster as *NGFR-*high, noting that both clusters contained cells from each duration of treatment. We then identified 44 resistant clones with at least five cells (as defined above) after four weeks of treatment with dabrafenib and trametinib. We then calculated the percentage of cells from each resistant clone that were in the *EGFR-* or *NGFR*-high cluster. We then followed these same 44 clones through to the 6.5 week treatment sample and identified whether a clone died (defined as no longer being identified as a resistance clone as defined above) and calculated the percentage of each surviving clone in each cluster. We then more broadly assess the survival of the clones by counting how many cells from each of the 44 clones were present in each of the three samples that had received a secondary round of treatment (dabrafenib and trametinib to dabrafenib and trametinib, dabrafenib and trametinib to CoCl_2_, and dabrafenib and trametinib to cisplatin).

### NGFR reporter analysis

We imaged the NGFR-mNG2(11) WM989 A6-G3 cell line^42^ on a Nikon Eclipse Ti2 microscope after combination treatment with dabrafenib and trametinib. We initially cultured the cell line in dabrafenib and trametinib-treated TU2% medium using the cell culture method noted above for about four weeks. At treatment day 32 and day 39, we imaged the cells on the aforementioned microscope. Prior to each scan, nuclei were transiently stained with Hoescht stain to visualize nuclei. We took images at 10x magnification of mNG2 fluorescence (600 ms in the green channel) and Hoescht fluorescence (200 ms in the blue channel). We chose this time series and these exposure times based on pilot experiments to minimize signs of cell death by phototoxicity and to enable the tracking of colonies.

For processing of these images, we converted ND2 images to TIFF format, and performed subsequent processing in ImageJ^56^. To improve NGFR-mNG2(11) signaling above background, we used the “Subtract Background” function, selecting the sliding paraboloid option and using a rolling ball radius of 1000 pixels. We then log scaled the pixels and added scale bars with a ratio of 1.29 μm per pixel. Images cropped from the large scan at each time point were contrast matched, but images between time points were not in order to properly visualize NGFR-mNG2(11) against background.

## Data Availability

All code and data used will be made available upon publication.

## Acknowledgements

We thank the members of the Shaffer Lab for input on the experiments and figures of the manuscript. We thank the Arjun Raj Lab at the University of Pennsylvania for the barcoding plasmid library. We thank ChatGPT for helping in paraphrasing and avoiding redundancy in our methods section. A.J.F., P.E.W. and S.M.S. recognize support from Grants for Faculty Mentoring Undergraduate Research (A.J.F. and S.M.S. 2021; P.E.W. and S.M.S. 2022). S.M.S acknowledges support from the NIH Director’s Early Independence Award DP5OD028144 and the Wistar/Penn Skin Cancer SPORE (P50 CA174523).

## Author Contributions

Conceptualization, D.L.S., A.J.F., and S.M.S.; Methodology, D.L.S., A.J.F., and S.M.S.; Software, D.L.S., A.J.F., and R.J.V.V.; Formal Analysis, D.L.S., A.J.F., and R.J.V.V.; Investigation, D.L.S., A.J.F., and P.E.W.; Data Curation, D.L.S., A.J.F., P.E.W., and R.J.V.V.; Writing - Original Draft, D.L.S., A.J.F., P.E.W., R. J.V.V., and S.M.S.; Writing - Review & Editing, D.L.S., A.J.F., P.E.W., R.J.V.V., and S.M.S.; Visualization, D.L.S., A.J.F., P.E.W., R.J.V.V., and S.M.S.; Funding Acquisition, S.M.S., A.J.F., and P.E.W; Supervision, S. M.S.

## Notes

### Competing Interest Statement

The authors have declared no competing interest.

## References

1. Nowell, P.C. (1976). The clonal evolution of tumor cell populations. Science 194, 23–28.

2. Jin, J., Wu, X., Yin, J., Li, M., Shen, J., Li, J., Zhao, Y., Zhao, Q., Wu, J., Wen, Q., et al. (2019). Identification of Genetic Mutations in Cancer: Challenge and Opportunity in the New Era of Targeted Therapy. Front. Oncol. 9, 263.

3. McGranahan, N., and Swanton, C. (2017). Clonal Heterogeneity and Tumor Evolution: Past, Present, and the Future. Cell 168, 613–628.

4. Marine, J.-C., Dawson, S.-J., and Dawson, M.A. (2020). Non-genetic mechanisms of therapeutic resistance in cancer. Nat. Rev. Cancer 20, 743–756.

5. Sharma, S.V., Lee, D.Y., Li, B., Quinlan, M.P., Takahashi, F., Maheswaran, S., McDermott, U., Azizian, N., Zou, L., Fischbach, M.A., et al. (2010). A chromatin-mediated reversible drug-tolerant state in cancer cell subpopulations. Cell 141, 69–80.

6. Knoechel, B., Roderick, J.E., Williamson, K.E., Zhu, J., Lohr, J.G., Cotton, M.J., Gillespie, S.M., Fernandez, D., Ku, M., Wang, H., et al. (2014). An epigenetic mechanism of resistance to targeted therapy in T cell acute lymphoblastic leukemia. Nat. Genet. 46, 364–370.

7. Spencer, S.L., Gaudet, S., Albeck, J.G., Burke, J.M., and Sorger, P.K. (2009). Non-genetic origins of cell-to-cell variability in TRAIL-induced apoptosis. Nature 459, 428–432.

8. Shaffer, S.M., Dunagin, M.C., Torborg, S.R., Torre, E.A., Emert, B., Krepler, C., Beqiri, M., Sproesser, K., Brafford, P.A., Xiao, M., et al. (2017). Rare cell variability and drug-induced reprogramming as a mode of cancer drug resistance. Nature 546, 431–435.

9. Goyal, Y., Dardani, I.P., Busch, G.T., Emert, B., Fingerman, D., Kaur, A., Jain, N., Mellis, I.A., Li, J., Kiani, K., et al. (2021). Pre-determined diversity in resistant fates emerges from homogenous cells after anti-cancer drug treatment. bioRxiv, 2021.12.08.471833. 10.1101/2021.12.08.471833.

10. Vander Velde, R., Yoon, N., Marusyk, V., Durmaz, A., Dhawan, A., Miroshnychenko, D., Lozano-Peral, D., Desai, B., Balynska, O., Poleszhuk, J., et al. (2020). Resistance to targeted therapies as a multifactorial,gradual adaptation to inhibitor specific selective pressures. Nat. Commun. 11, 2393.

11. Harmange, G., Reyes Hueros, R.A., Schaff, D., Emert, B., Saint-Antoine, M., Nellore, S., Fane, M.E., Alicea, G.M., Weeraratna, A.T., Singh, A., et al. (2022). Disrupting cellular memory to overcome drug resistance. bioRxiv, 2022.06.16.496161. 10.1101/2022.06.16.496161.

12. Beaumont, K.A., Hill, D.S., Daignault, S.M., Lui, G.Y.L., Sharp, D.M., Gabrielli, B., Weninger, W., and Haass, N.K. (2016). Cell Cycle Phase-Specific Drug Resistance as an Escape Mechanism of Melanoma Cells. J. Invest. Dermatol. 136, 1479–1489.

13. Mishra, A., Wang, J., Shiozawa, Y., McGee, S., Kim, J., Jung, Y., Joseph, J., Berry, J.E., Havens, A., Pienta, K.J., et al. (2012). Hypoxia Stabilizes GAS6/Axl Signaling in Metastatic Prostate Cancer. Molecular Cancer Research 10, 703–712. 10.1158/1541-7786.mcr-11-0569.

14. Widmer, D.S., Hoek, K.S., Cheng, P.F., Eichhoff, O.M., Biedermann, T., Raaijmakers, M.I.G., Hemmi, S., Dummer, R., and Levesque, M.P. (2013). Hypoxia contributes to melanoma heterogeneity by triggering HIF1α-dependent phenotype switching. J. Invest. Dermatol. 133, 2436–2443.

15. Ravindran Menon, D., Das, S., Krepler, C., Vultur, A., Rinner, B., Schauer, S., Kashofer, K., Wagner, K., Zhang, G., Bonyadi Rad, E., et al. (2015). A stress-induced early innate response causes multidrug tolerance in melanoma. Oncogene 34, 4448–4459.

16. Erdmann, S., Seidel, D., Jahnke, H.-G., Eichler, M., Simon, J.-C., and Robitzki, A.A. (2019). Induced cross-resistance of BRAFV600E melanoma cells to standard chemotherapeutic dacarbazine after chronic PLX4032 treatment. Sci. Rep. 9, 30.

17. Stordal, B., Pavlakis, N., and Davey, R. (2007). Oxaliplatin for the treatment of cisplatin-resistant cancer: a systematic review. Cancer Treat. Rev. 33, 347–357.

18. Fojo, A., Hamilton, T.C., Young, R.C., and Ozols, R.F. (1987). Multidrug resistance in ovarian cancer. Cancer 60, 2075–2080.

19. Menzies, A.M., Long, G.V., and Murali, R. (2012). Dabrafenib and its potential for the treatment of metastatic melanoma. Drug Des. Devel. Ther. 6, 391–405.

20. Lugowska, I., Koseła-Paterczyk, H., Kozak, K., and Rutkowski, P. (2015). Trametinib: a MEK inhibitor for management of metastatic melanoma. Onco. Targets. Ther. 8, 2251–2259.

21. Piret, J.-P., Mottet, D., Raes, M., and Michiels, C. (2002). CoCl2, a chemical inducer of hypoxia-inducible factor-1, and hypoxia reduce apoptotic cell death in hepatoma cell line HepG2. Ann. N. Y. Acad. Sci. 973, 443–447.

22. Wu, D., and Yotnda, P. (2011). Induction and testing of hypoxia in cell culture. J. Vis. Exp. 10.3791/2899.

23. Sherman, S.E., and Lippard, S.J. (1987). Structural aspects of platinum anticancer drug interactions with DNA. Chem. Rev. 87, 1153–1181.

24. Biddy, B.A., Kong, W., Kamimoto, K., Guo, C., Waye, S.E., Sun, T., and Morris, S.A. (2018). Single-cell mapping of lineage and identity in direct reprogramming. Nature 564, 219–224.

25. Weinreb, C., Rodriguez-Fraticelli, A., Camargo, F.D., and Klein, A.M. (2020). Lineage tracing on transcriptional landscapes links state to fate during differentiation. Science 367. 10.1126/science.aaw3381.

26. Emert, B.L., Cote, C.J., Torre, E.A., Dardani, I.P., Jiang, C.L., Jain, N., Shaffer, S.M., and Raj, A. (2021). Variability within rare cell states enables multiple paths toward drug resistance. Nat. Biotechnol. 39, 865–876.

27. França, G.S., Baron, M., Pour, M., King, B.R., Rao, A., Misirlioglu, S., Barkley, D., Dolgalev, I., Ho-Tang, K., Avital, G., et al. (2022). Drug-induced adaptation along a resistance continuum in cancer cells. bioRxiv, 2022.06.21.496830. 10.1101/2022.06.21.496830.

28. Rose-John, S. (2020). Interleukin-6 signalling in health and disease. F1000Res. 9. 10.12688/f1000research.26058.1.

29. Wang, Y., Niu, X.L., Qu, Y., Wu, J., Zhu, Y.Q., Sun, W.J., and Li, L.Z. (2010). Autocrine production of interleukin-6 confers cisplatin and paclitaxel resistance in ovarian cancer cells. Cancer Lett. 295, 110–123.

30. Zhao, K., Lu, Y., Chen, Y., Cheng, J., and Zhang, W. (2020). Transcripts 202 and 205 of IL-6 confer resistance to Vemurafenib by reactivating the MAPK pathway in BRAF(V600E) mutant melanoma cells. Exp. Cell Res. 390, 111942.

31. Masjedi, A., Hashemi, V., Hojjat-Farsangi, M., Ghalamfarsa, G., Azizi, G., Yousefi, M., and Jadidi-Niaragh, F. (2018). The significant role of interleukin-6 and its signaling pathway in the immunopathogenesis and treatment of breast cancer. Biomed. Pharmacother. 108, 1415–1424.

32. Niu, N., Yao, J., Bast, R.C., Sood, A.K., and Liu, J. (2021). IL-6 promotes drug resistance through formation of polyploid giant cancer cells and stromal fibroblast reprogramming. Oncogenesis 10, 65.

33. Wu, T., Hu, E., Xu, S., Chen, M., Guo, P., Dai, Z., Feng, T., Zhou, L., Tang, W., Zhan, L., et al. (2021). clusterProfiler 4.0: A universal enrichment tool for interpreting omics data. Innovation (Camb) 2, 100141.

34. Fedorenko, I.V., Abel, E.V., Koomen, J.M., Fang, B., Wood, E.R., Chen, Y.A., Fisher, K.J., Iyengar, S., Dahlman, K.B., Wargo, J.A., et al. (2016). Fibronectin induction abrogates the BRAF inhibitor response of BRAF V600E/PTEN-null melanoma cells. Oncogene 35, 1225–1235.

35. Marusak, C., Thakur, V., Li, Y., Freitas, J.T., Zmina, P.M., Thakur, V.S., Chang, M., Gao, M., Tan, J., Xiao, M., et al. (2020). Targeting Extracellular Matrix Remodeling Restores BRAF Inhibitor Sensitivity in BRAFi-resistant Melanoma. Clin. Cancer Res. 26, 6039–6050.

36. Straussman, R., Morikawa, T., Shee, K., Barzily-Rokni, M., Qian, Z.R., Du, J., Davis, A., Mongare, M.M., Gould, J., Frederick, D.T., et al. (2012). Tumour micro-environment elicits innate resistance to RAF inhibitors through HGF secretion. Nature 487, 500–504.

37. Czarnecka, A.M., Bartnik, E., Fiedorowicz, M., and Rutkowski, P. (2020). Targeted Therapy in Melanoma and Mechanisms of Resistance. Int. J. Mol. Sci. 21. 10.3390/ijms21134576.

38. MacKay, C., Carroll, E., Ibrahim, A.F.M., Garg, A., Inman, G.J., Hay, R.T., and Alpi, A.F. (2014). E3 ubiquitin ligase HOIP attenuates apoptotic cell death induced by cisplatin. Cancer Res. 74, 2246–2257.

39. Ko, T., and Li, S. (2019). Genome-wide screening identifies novel genes and biological processes implicated in cisplatin resistance. FASEB J. 33, 7143–7154.

40. Ashton, T.M., McKenna, W.G., Kunz-Schughart, L.A., and Higgins, G.S. (2018). Oxidative Phosphorylation as an Emerging Target in Cancer Therapy. Clin. Cancer Res. 24, 2482–2490.

41. Li, L.-D., Sun, H.-F., Liu, X.-X., Gao, S.-P., Jiang, H.-L., Hu, X., and Jin, W. (2015). Down-Regulation of NDUFB9 Promotes Breast Cancer Cell Proliferation, Metastasis by Mediating Mitochondrial Metabolism. PLoS One 10, e0144441.

42. Shaffer, S.M., Emert, B.L., Reyes Hueros, R.A., Cote, C., Harmange, G., Schaff, D.L., Sizemore, A.E., Gupte, R., Torre, E., Singh, A., et al. (2020). Memory Sequencing Reveals Heritable Single-Cell Gene Expression Programs Associated with Distinct Cellular Behaviors. Cell 182, 947–959.e17.

43. Sartor, O., and Gillessen, S. (2014). Treatment sequencing in metastatic castrate-resistant prostate cancer. Asian J. Androl. 16, 426–431.

44. Modest, D.P., Pant, S., and Sartore-Bianchi, A. (2019). Treatment sequencing in metastatic colorectal cancer. Eur. J. Cancer 109, 70–83.

45. Temraz, S., Mukherji, D., and Shamseddine, A. (2014). Sequencing of treatment in metastatic colorectal cancer: where to fit the target. World J. Gastroenterol. 20, 1993–2004.

46. Johnson, D.B., Pectasides, E., Feld, E., Ye, F., Zhao, S., Johnpulle, R., Merritt, R., McDermott, D.F., Puzanov, I., Lawrence, D., et al. (2017). Sequencing Treatment in BRAF V600 Mutant Melanoma:Anti-PD-1 Before and After BRAF Inhibition. J. Immunother. 40, 31–35.

47. Girotti, M.R., Gremel, G., Lee, R., Galvani, E., Rothwell, D., Viros, A., Mandal, A.K., Lim, K.H.J., Saturno, G., Furney, S.J., et al. (2016). Application of Sequencing, Liquid Biopsies, and Patient-Derived Xenografts for Personalized Medicine in Melanoma. Cancer Discov. 6, 286–299.

48. Buck, S.A.J., Koolen, S.L.W., Mathijssen, R.H.J., de Wit, R., and van Soest, R.J. (2021). Cross-resistance and drug sequence in prostate cancer. Drug Resist. Updat. 56, 100761.

49. Vander Velde, R., Shaffer, S., and Marusyk, A. (2022). Integrating mutational and nonmutational mechanisms of acquired therapy resistance within the Darwinian paradigm. Trends Cancer Res. 8, 456–466.

50. Roesch, A., Vultur, A., Bogeski, I., Wang, H., Zimmermann, K.M., Speicher, D., Körbel, C., Laschke, M.W., Gimotty, P.A., Philipp, S.E., et al. (2013). Overcoming intrinsic multidrug resistance in melanoma by blocking the mitochondrial respiratory chain of slow-cycling JARID1B(high) cells. Cancer Cell 23, 811–825.

51. Roesch, A., Fukunaga-Kalabis, M., Schmidt, E.C., Zabierowski, S.E., Brafford, P.A., Vultur, A., Basu, D., Gimotty, P., Vogt, T., and Herlyn, M. (2010). A temporarily distinct subpopulation of slow-cycling melanoma cells is required for continuous tumor growth. Cell 141, 583–594.

52. Hao, Y., Hao, S., Andersen-Nissen, E., Mauck, W.M., 3rd, Zheng, S., Butler, A., Lee, M.J., Wilk, A.J., Darby, C., Zager, M., et al. (2021). Integrated analysis of multimodal single-cell data. Cell 184, 3573–3587.e29.

53. Subramanian, A., Tamayo, P., Mootha, V.K., Mukherjee, S., Ebert, B.L., Gillette, M.A., Paulovich, A., Pomeroy, S.L., Golub, T.R., Lander, E.S., et al. (2005). Gene set enrichment analysis: a knowledge-based approach for interpreting genome-wide expression profiles. Proc. Natl. Acad. Sci. U. S. A. 102, 15545–15550.

54. Liberzon, A., Birger, C., Thorvaldsdóttir, H., Ghandi, M., Mesirov, J.P., and Tamayo, P. (2015). The Molecular Signatures Database (MSigDB) hallmark gene set collection. Cell Syst 1, 417–425.

55. Korotkevich, G., Sukhov, V., Budin, N., Shpak, B., Artyomov, M.N., and Sergushichev, A. (2021). Fast gene set enrichment analysis. bioRxiv, 060012. 10.1101/060012.

56. Schneider, C.A., Rasband, W.S., and Eliceiri, K.W. (2012). NIH Image to Image J: 25 years of image analysis. Nat. Methods 9, 671–675.

