## Supplemental Figures for "Clonal differences underlie variable responses to sequential and prolonged treatment"

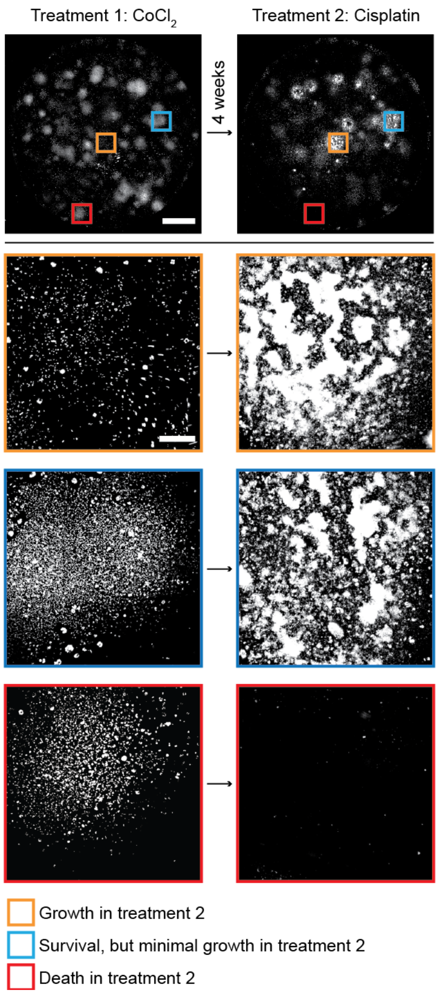


**Supplemental Figure 1: Cancer cell clones exhibit variable responses to first and second treatments.** WM989 BRAF V600E mutant melanoma cells with a nuclear GFP tag imaged after four weeks in $CoCl_2$ followed by four weeks in cisplatin (top). Below are cropped images from the whole well scans showing clones that grew (orange), survived with low growth (blue), or mostly died (red) in the second treatment. Cisplatin resistant cells tend to grow on top of each other causing cropped images to look out of focus or overexposed. Scale bars in whole well scans are 5 mm while scale bars in cropped scans are 500 µm.

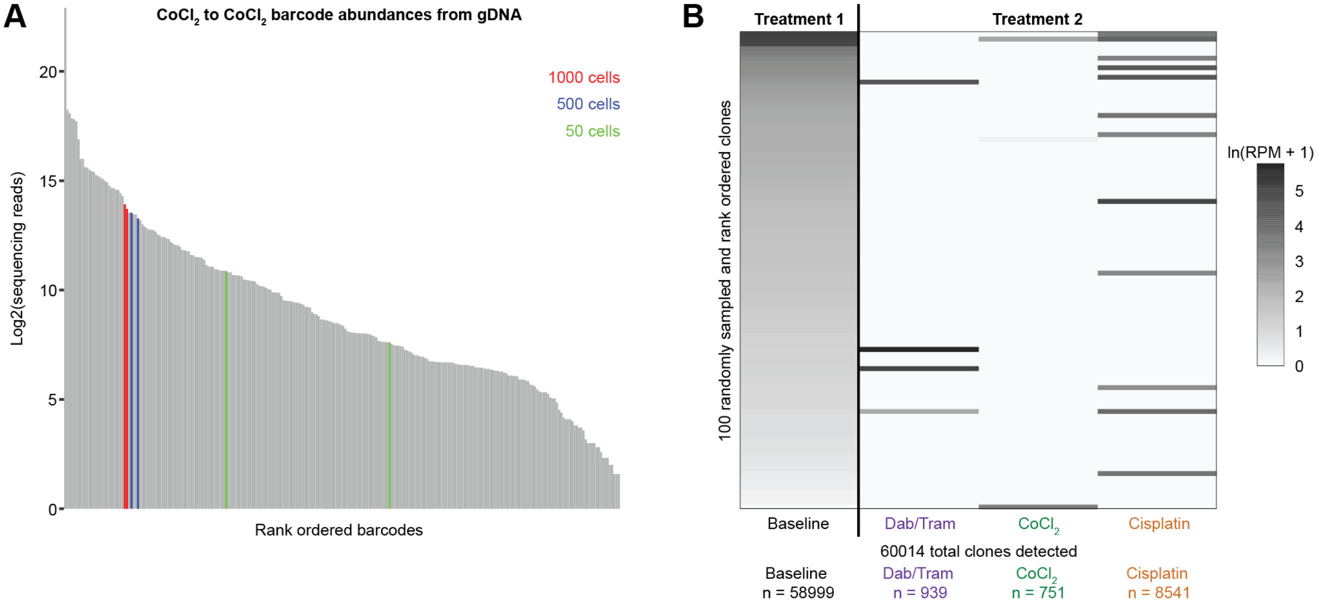


**Supplemental Figure 2: Clonal barcode reads are a quantitative readout for cell number. A)** Sequencing reads of DNA barcodes from gDNA using the $CoCl_2$ to $CoCl_2$ condition as an example. Displayed are the $log_2$(# of sequencing reads) with the ladder barcodes highlighted in red (1000 cells), blue (500 cells), and green (50 cells). Only barcodes with at least two reads are displayed. **B)** Heatmaps showing 100 randomly sampled and rank-ordered clones immediately prior to treatment being applied (baseline) followed by their abundance after initial treatment with a combination of dabrafenib and trametinib (Dab/Tram), $CoCl_2$, and cisplatin. Individual clones are colored by the ln(Reads Per Million (RPM) + 1) using data from sequencing clonal barcodes from gDNA of surviving cells. It should be noted that there are clones whose abundance were below the threshold of detection in the pretreatment sample, but were detected after the first round of treatment. The total number of unique clones that were detected before or after initial treatment is displayed below the heatmap, followed by the number of clones detected after each treatment.

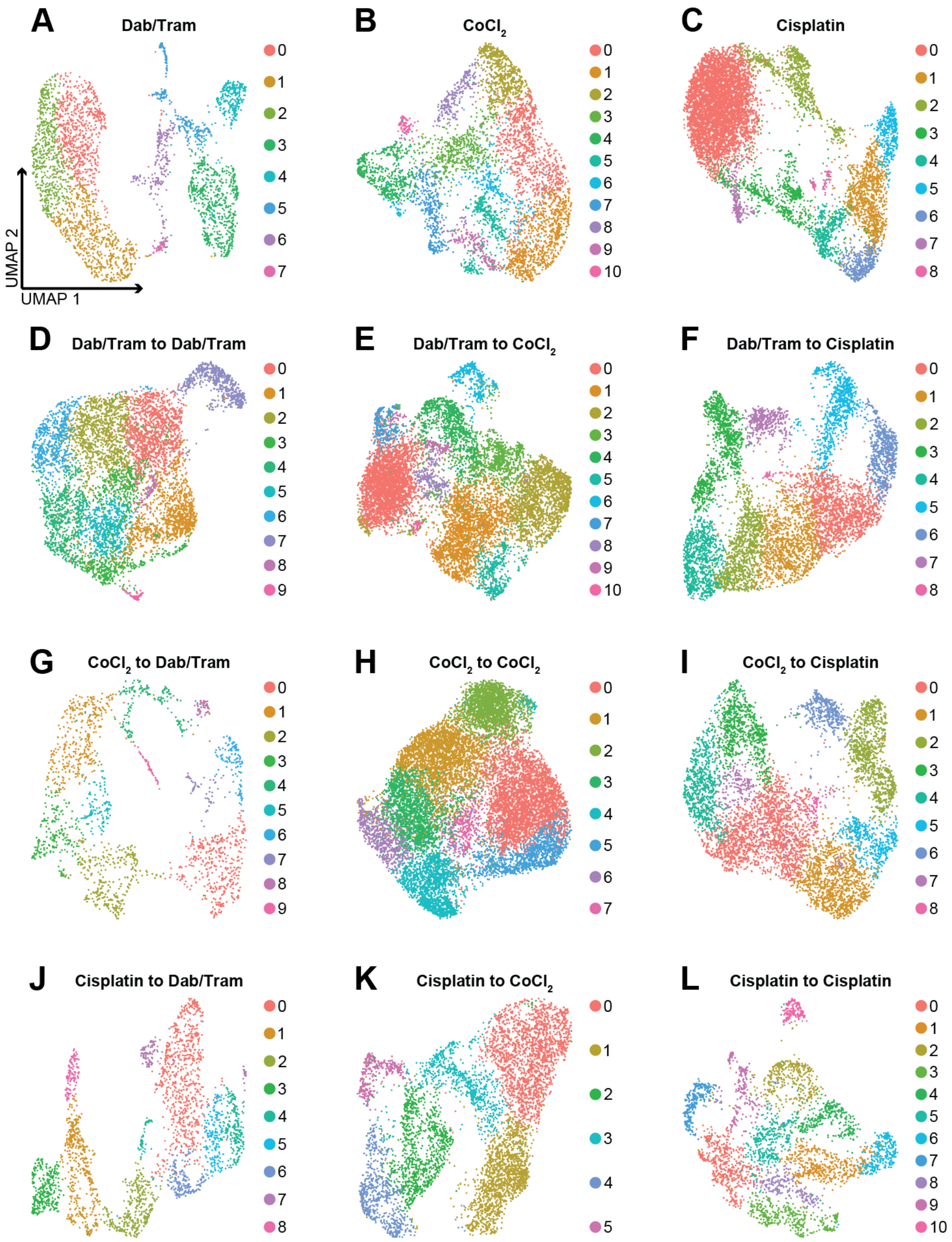


**Supplemental Figure 3: UMAPs of each condition individually.** UMAP projections of **A)** dabrafenib and trametinib (Dab/Tram, 3203 cells), **B)** $CoCl_2$ (4823 cells), **C)** cisplatin (8303 cells), **D)** Dab/Tram to Dab/Tram (6906 cells), **E)** Dab/Tram to $CoCl_2$ (10184 cells), **F)** Dab/Tram to cisplatin (7070 cells), **G)** $CoCl_2$ to Dab/Tram (1773 cells), **H)** $CoCl_2$ to $CoCl_2$ (13951 cells), **I)** $CoCl_2$ to cisplatin (6459 cells), **J)** cisplatin to Dab/Tram (2615 cells), **K)** cisplatin to $CoCl_2$ (5715 cells), and **L)** cisplatin to cisplatin (3323 cells) treated cells. The UMAP projections are colored based on their default Seurat[52] clustering.

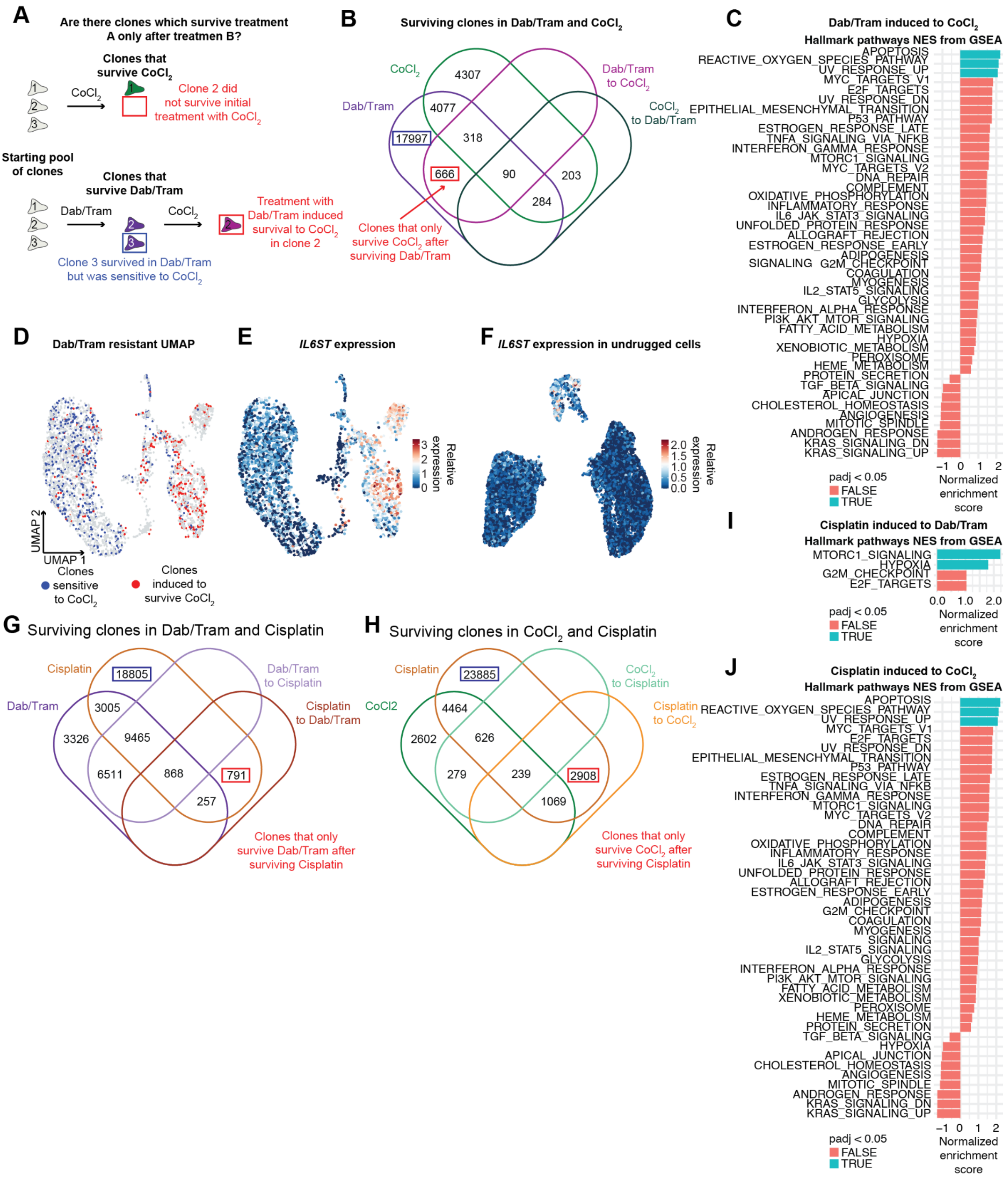


**Supplemental Figure 4: Initial treatment induces resistance to a second therapy in previously sensitive clones.** We asked the question of whether there are clones that survive treatment A only after treatment B. Panel **A** shows a schematic for identifying clones that are only able to survive a treatment after being treated with a different treatment (induced resistance), using $CoCl_2$ and a combination of dabrafenib and trametinib (Dab/Tram) as an example. Beginning with the same pool of starting clones, clone two does not survive initial treatment with $CoCl_2$. However, clone two does survive $CoCl_2$ if it had previously survived treatment with Dab/Tram. This is contrasted with clone three that survived Dab/Tram, but was unable to survive $CoCl_2$ treatment. We only showed venn diagrams for comparisons that yielded differentially enriched hallmark pathways by GSEA in induced clones[53–55]. **B)** Venn diagram comparing clones that survived Dab/Tram, $CoCl_2$, Dab/Tram to $CoCl_2$, and $CoCl_2$ to Dab/Tram. Boxed in red are the 666 clones that only ever survived $CoCl_2$ after previous treatment with Dab/Tram. Boxed in blue are the 17997 clones that survived Dab/Tram but were still sensitive to $CoCl_2$. **C)** We identified genes differentially expressed between clones boxed in blue and red in **B** to find pathways enriched and depleted in clones that were induced to survive $CoCl2_2$ by Dab/Tram. All pathways displayed were significantly enriched ($p < 0.05$), the blue bars indicate that the adjusted p value was also < 0.05. **D)** Dab/Tram resistant UMAP with cells colored as red if they are a clone with induced survival to $CoCl_2$ and blue if they are sensitive to $CoCl_2$. The blue cells clustered largely on the left and the red largely on the right. **E)** Relative gene expression of *IL6ST*. High expression aligns with the same area of the UMAP as induced clones in **D**. **F)** Relative *IL6ST* expression in untreated cells from Harmange et al.[11]. **G)** Venn diagram comparing clones that survived Dab/Tram, cisplatin, Dab/Tram to cisplatin, and cisplatin to Dab/Tram. Boxed in red are the 791 clones that only ever survived Dab/Tram after previous treatment with cisplatin. Boxed in blue are the 18805 clones that survived cisplatin but were still sensitive to Dab/Tram. **H)** Venn diagram comparing clones that survived $CoCl_2$, cisplatin, $CoCl_2$ to cisplatin, and cisplatin to $CoCl_2$. Boxed in red are the 2908 clones that only ever survived Dab/Tram after previous treatment with cisplatin. Boxed in blue are the 23885 clones that survived cisplatin but were still sensitive to Dab/Tram. **I)** Differentially expressed pathways in clones with resistance to Dab/Tram induced by cisplatin. All pathways displayed were significantly enriched ($p < 0.05$), the blue bars indicate that the adjusted p value was also < 0.05. **J)** Differentially expressed pathways in clones with resistance to $CoCl_2$ induced by cisplatin. All pathways displayed were significantly enriched ($p < 0.05$), the blue bars indicate that the adjusted p value was also < 0.05.

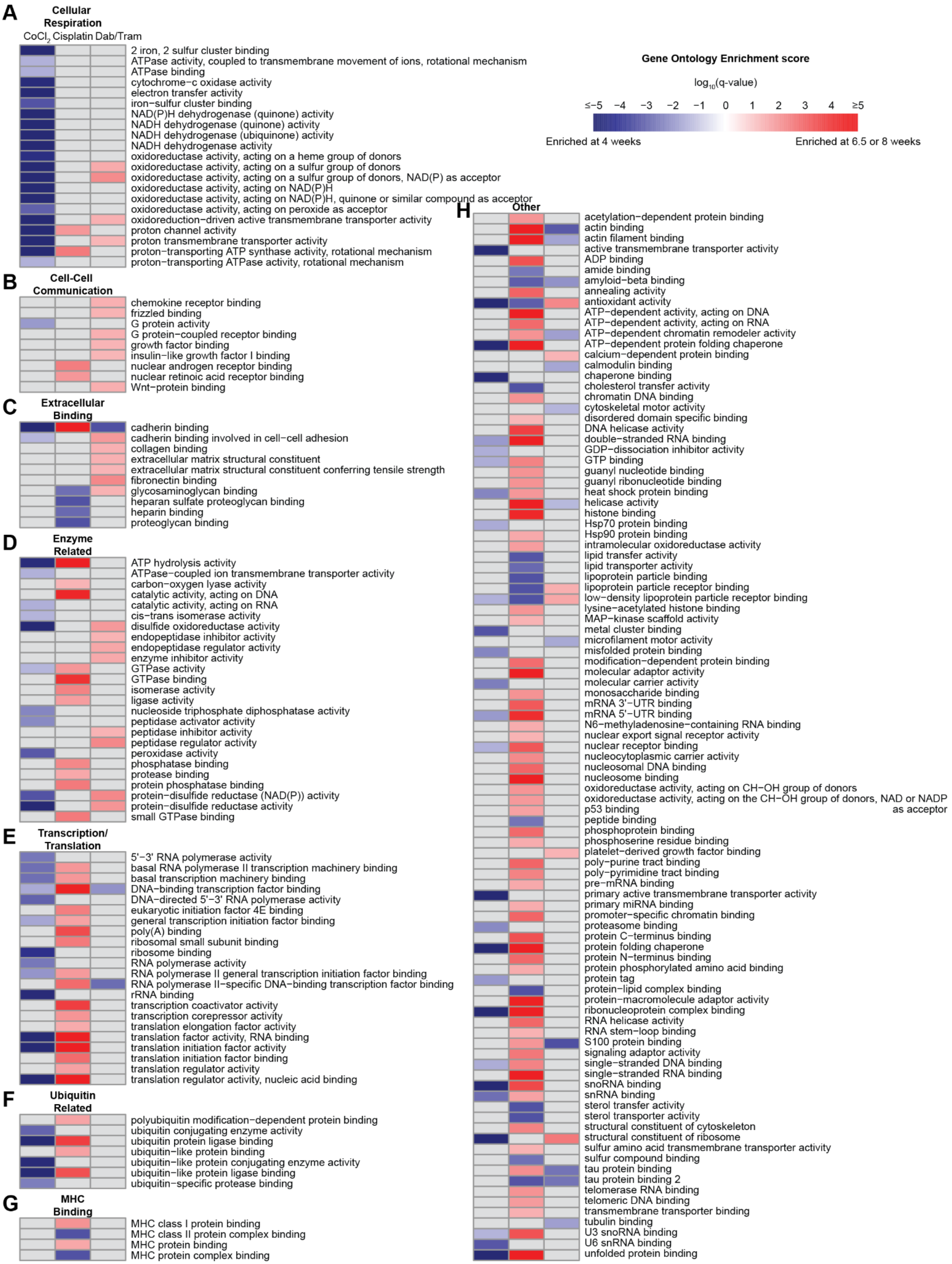
A
Cellular Respiration
CoCl2 Cisplatin Dab/Tram
2 iron, 2 sulfur cluster binding
ATPase activity, coupled to transmembrane movement of ions, rotational mechanism
ATPase binding
cytochrome−c oxidase activity
electron transfer activity
iron−sulfur cluster binding
NAD(P)H dehydrogenase (quinone) activity
NADH dehydrogenase (quinone) activity
NADH dehydrogenase (ubiquinone) activity
NADH dehydrogenase activity
oxidoreductase activity, acting on a heme group of donors
oxidoreductase activity, acting on a sulfur group of donors
oxidoreductase activity, acting on a sulfur group of donors, NAD(P) as acceptor
oxidoreductase activity, acting on NAD(P)H
oxidoreductase activity, acting on NAD(P)H, quinone or similar compound as acceptor
oxidoreductase activity, acting on peroxide as acceptor
oxidoreduction−driven active transmembrane transporter activity
proton channel activity
proton transmembrane transporter activity
proton−transporting ATP synthase activity, rotational mechanism
proton−transporting ATPase activity, rotational mechanism
B
Cell-Cell Communication
chemokine receptor binding
frizzled binding
G protein activity
G protein−coupled receptor binding
growth factor binding
insulin−like growth factor I binding
nuclear androgen receptor binding
nuclear retinoic acid receptor binding
Wnt−protein binding
C
Extracellular Binding
cadherin binding
cadherin binding involved in cell−cell adhesion
collagen binding
extracellular matrix structural constituent
extracellular matrix structural constituent conferring tensile strength
fibronectin binding
glycosaminoglycan binding
heparan sulfate proteoglycan binding
heparin binding
proteoglycan binding
D
Enzyme Related
ATP hydrolysis activity
ATPase−coupled ion transmembrane transporter activity
carbon−oxygen lyase activity
catalytic activity, acting on DNA
catalytic activity, acting on RNA
cis−trans isomerase activity
disulfide oxidoreductase activity
endopeptidase inhibitor activity
endopeptidase regulator activity
enzyme inhibitor activity
GTPase activity
GTPase binding
isomerase activity
ligase activity
nucleoside triphosphate diphosphatase activity
peptidase activator activity
peptidase inhibitor activity
peptidase regulator activity
peroxidase activity
phosphatase binding
protease binding
protein phosphatase binding
protein−disulfide reductase (NAD(P)) activity
protein−disulfide reductase activity
small GTPase binding
E
Transcription/ Translation
5'−3' RNA polymerase activity
basal RNA polymerase II transcription machinery binding
basal transcription machinery binding
DNA−binding transcription factor binding
DNA−directed 5'−3' RNA polymerase activity
eukaryotic initiation factor 4E binding
general transcription initiation factor binding
poly(A) binding
ribosomal small subunit binding
ribosome binding
RNA polymerase activity
RNA polymerase II general transcription initiation factor binding
RNA polymerase II−specific DNA−binding transcription factor binding
rRNA binding
transcription coactivator activity
transcription corepressor activity
translation elongation factor activity
translation factor activity, RNA binding
translation initiation factor activity
translation initiation factor binding
translation regulator activity
translation regulator activity, nucleic acid binding
F
Ubiquitin Related
polyubiquitin modification−dependent protein binding
ubiquitin conjugating enzyme activity
ubiquitin protein ligase binding
ubiquitin−like protein binding
ubiquitin−like protein conjugating enzyme activity
ubiquitin−like protein ligase binding
ubiquitin−specific protease binding
G
MHC Binding
MHC class I protein binding
MHC class II protein complex binding
MHC protein binding
MHC protein complex binding
Gene Ontology Enrichment score
log10(q-value)
≤−5 −4 −3 −2 −1 0 1 2 3 4 ≥5
Enriched at 4 weeks
Enriched at 6.5 or 8 weeks
H
Other
acetylation−dependent protein binding
actin binding
actin filament binding
active transmembrane transporter activity
ADP binding
amide binding
amyloid−beta binding
annealing activity
antioxidant activity
ATP−dependent activity, acting on DNA
ATP−dependent activity, acting on RNA
ATP−dependent chromatin remodeler activity
ATP−dependent protein folding chaperone
calcium−dependent protein binding
calmodulin binding
chaperone binding
cholesterol transfer activity
chromatin DNA binding
cytoskeletal motor activity
disordered domain specific binding
DNA helicase activity
double−stranded RNA binding
GDP−dissociation inhibitor activity
GTP binding
guanyl nucleotide binding
guanyl ribonucleotide binding
heat shock protein binding
helicase activity
histone binding
Hsp70 protein binding
Hsp90 protein binding
intramolecular oxidoreductase activity
lipid transfer activity
lipid transporter activity
lipoprotein particle binding
lipoprotein particle receptor binding
low−density lipoprotein particle receptor binding
lysine−acetylated histone binding
MAP−kinase scaffold activity
metal cluster binding
microfilament motor activity
misfolded protein binding
modification−dependent protein binding
molecular adaptor activity
molecular carrier activity
monosaccharide binding
mRNA 3'−UTR binding
mRNA 5'−UTR binding
N6−methyladenosine−containing RNA binding
nuclear export signal receptor activity
nuclear receptor binding
nucleocytoplasmic carrier activity
nucleosomal DNA binding
nucleosome binding
oxidoreductase activity, acting on CH−OH group of donors
oxidoreductase activity, acting on the CH−OH group of donors, NAD or NADP as acceptor
p53 binding
peptide binding
phosphoprotein binding
phosphoserine residue binding
platelet−derived growth factor binding
poly−purine tract binding
poly−pyrimidine tract binding
pre−mRNA binding
primary active transmembrane transporter activity
primary miRNA binding
promoter−specific chromatin binding
proteasome binding
protein C−terminus binding
protein folding chaperone
protein N−terminus binding
protein phosphorylated amino acid binding
protein tag
protein−lipid complex binding
protein−macromolecule adaptor activity
ribonucleoprotein complex binding
RNA helicase activity
RNA stem−loop binding
S100 protein binding
signaling adaptor activity
single−stranded DNA binding
single−stranded RNA binding
snoRNA binding
snRNA binding
sterol transfer activity
sterol transporter activity
structural constituent of cytoskeleton
structural constituent of ribosome
sulfur amino acid transmembrane transporter activity
sulfur compound binding
tau protein binding
tau protein binding 2
telomerase RNA binding
telomeric DNA binding
transmembrane transporter binding
tubulin binding
U3 snoRNA binding
U6 snRNA binding
unfolded protein binding

**Supplemental Figure 5: Gene Ontology comparing cells in the same agent after first and second treatment.** We compared cells from the same agent after the first and second round of treatment, identifying differentially expressed genes between the two. We then identified GO terms enriched after either the first round of treatment (four weeks, blue) or after the second round of treatment (6.5 or eight weeks, red) based on the $\log_{10}$(q-value) and $-\log_{10}$(q-value) respectively. GO terms that were not significantly enriched, or failed other thresholds, in either direction for a condition are colored in gray. Subpanels **A-G** show the same pathways in the same order as in Figure 5A, showing the pathways that make up each category. **H)** The “other” category includes all of the pathways that were significant in at least one of the three comparisons that we did not group into one of the seven other categories. Column labels indicate the agent from which cells after the first and second round of treatment were compared.

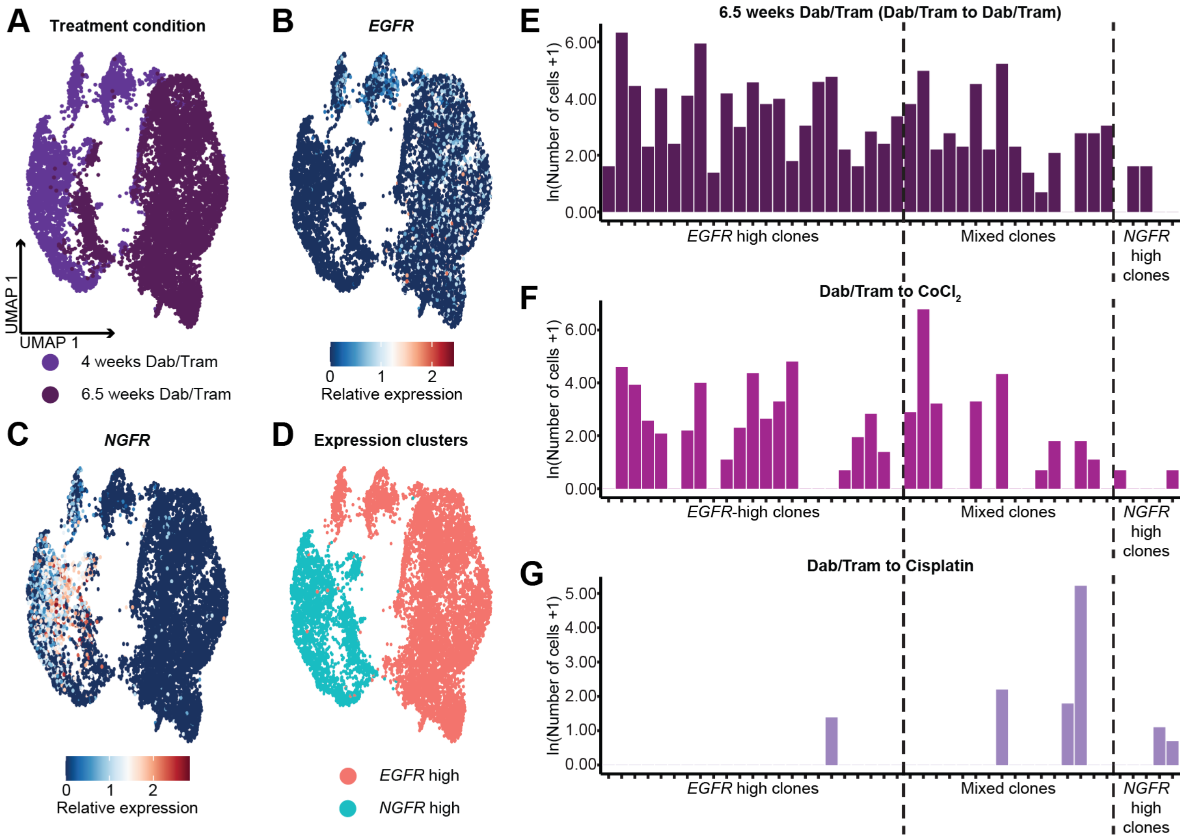


**Supplemental Figure 6: Dabrafenib and trametinib treated cells enter either an *EGFR*- or *NGFR*-high state. A)** UMAP projection of cells that have only been treated with dabrafenib and trametinib (Dab/Tram) after first (four weeks) and second (6.5 weeks) rounds of treatment. The cells cluster predominantly by condition, but also have overlap between conditions. Relative expression of **B)** *EGFR* or **C)** *NGFR*. Cells expressing these markers are largely mutually exclusive. **D)** We constrained Seurat clustering to only allow two total clusters[52]. We denoted these clusters as *EGFR*-high and *NGFR*-high based on their correlation with the markers. We identified clones with at least five cells after initial Dab/Tram treatment and classified them as *EGFR*-high, mixed clones, and *NGFR*-high. We then measured the number of sequenced single-cells from these clones after secondary treatment with **E)** Dab/Tram, **F)** $CoCl_2$, **G)** and cisplatin treatment, displayed as ln(# of cells + 1).
